# Genome assembly and characterization of a complex zfBED-NLR gene-containing disease resistance locus in Carolina Gold Select rice with Nanopore sequencing

**DOI:** 10.1101/675678

**Authors:** Andrew C. Read, Matthew J. Moscou, Aleksey V. Zimin, Geo Pertea, Rachel S. Meyer, Michael D. Purugganan, Jan E. Leach, Lindsay R. Triplett, Steven L. Salzberg, Adam J. Bogdanove

## Abstract

**Background:** Long-read sequencing facilitates assembly of complex genomic regions. In plants, loci containing nucleotide-binding, leucine-rich repeat (NLR) disease resistance genes are an important example of such regions. NLR genes make up one of the largest gene families in plants and are often clustered, evolving via duplication, contraction, and transposition. We recently mapped the *Xo1* locus for resistance to bacterial blight and bacterial leaf streak, found in the American heirloom rice variety Carolina Gold Select, to a region that in the Nipponbare reference genome is rich in NLR genes.

**Results:** Toward identification of the *Xo1* gene, we combined Nanopore and Illumina reads to generate a high-quality genome assembly for Carolina Gold Select. We identified 529 full or partial NLR genes and discovered, relative to the reference, an expansion of NLR genes at the *Xo1* locus. One NLR gene at *Xo1* has high sequence similarity to the cloned, functionally similar *Xa1* gene. Both harbor an integrated zfBED domain and near-identical, tandem, C-terminal repeats. Across diverse Oryzeae, we identified two sub-clades of such NLR genes, varying in the presence of the zfBED domain and the number of repeats.

**Conclusions:** Whole genome sequencing combining Nanopore and Illumina reads effectively resolves NLR gene loci, providing context as well as content. Our identification of an *Xo1* candidate is an important step toward mechanistic characterization, including the role(s) of the zfBED domain. Further, the Carolina Gold Select genome assembly will facilitate identification and exploitation of other useful traits in this historically important rice variety.

## BACKGROUND

Recent advances in sequencing technology enable the assembly of complex genomic loci by generating read lengths long enough to resolve repetitive regions [1]. Repetitive regions are often hotspots of recombination and other genomic changes, but difficulties assembling them mean that they often remain as incomplete gaps for many years after a genome’s initial draft assembly. For example, the centromeres and telomeres remain unsequenced for nearly all plant and animal genomes today. The most straightforward way to span lengthy or complex repeats is to generate single reads that are longer than the repeats themselves, so that repeats can be placed in the correct genomic location. When repeats occur in tandem arrays, reads need to be longer than the entire array if one is to accurately determine the number of repeat copies that the array contains. One of the most promising current technologies for resolving complex repeats is nanopore-based sequencing from Oxford Nanopore Technologies (“Nanopore”). Nanopore instruments pass DNA through a pore, monitor the change in electrical current across the pore, and convert the resulting signal into DNA sequence with ever-improving basecallers. Read-lengths are limited only by input DNA lengths, and validated reads as long as 2,272,580 bases have been reported [2]. Nanopore sequencing has been used for various applications, including genome sequencing of *Arabidopsis* and a wild tomato relative [3, 4], resolving complex T-DNA insertions [5], and disease resistance gene enrichment sequencing [6].

Plant disease resistance loci represent an important example of complex portions of a genome that can be challenging to characterize in context using short-read sequencing. These loci often contain clusters of nucleotide binding leucine-rich repeat (NLR) protein genes. NLR proteins are structurally modular, typically containing an N-terminal coiled-coil domain or a Toll/interleukin-1 receptor (TIR) domain, a conserved nucleotide binding domain (NB-ARC), and a C-terminal region comprising a variable number of leucine-rich repeats (LRRs). The NLR gene family is one of the largest and most diverse in plants [7, 8], with 95, 151, and 458 members reported in maize, *Arabidopsis*, and rice, respectively [9, 10]. Fifty-one percent of the rice NLR genes occur in 44 clusters in the genome [11]. Plants lack an adaptive immune system, and it has been theorized that this clustering provides plants an arsenal of resistance genes that can rapidly evolve, through duplication and recombination, to respond to dynamic pathogen populations [12–15]. Indeed, the structure and content of NLR loci is variable, even in closely related cultivars. Among plant populations, NLR genes account for the majority of copy-number and presence/absence polymorphisms [16–20]. Adding to the complexity of NLR genes, and the challenge of their sequence assembly, is the recent observation that approximately 10% of NLR genes encode additional, non-canonical, integrated domains (IDs) that may act as decoys, have roles in oligomerization or downstream signaling [21, 22], or serve other functions. Analysis of closely related species has shown that these IDs appear to be modular, with independent integrations occurring in diverse NLR genes over evolutionary time [23].

In this study, we sought to delineate NLR gene content at a disease resistance locus, *Xo1*, which we identified in 2016 in rice variety Carolina Gold Select [24], by using Nanopore long-reads combined with Illumina short-reads to generate a high quality, whole genome assembly. Carolina Gold Select is a purified line of Carolina Gold, a long-grain variety known for its distinctive gold hull and nutty flavor. Carolina Gold was the dominant variety grown in colonial America and is a breeding ancestor of modern US varieties [25]. Genotyping and draft genome sequencing confirmed it to be in the tropical Japonica clade [26, 27], but have not been sufficient to resolve loci associated with important disease resistance phenotypes in this variety, such as *Xo1*. *Xo1* protects against two important bacterial diseases, bacterial leaf streak (BLS) and bacterial blight (BB), caused by *Xanthomonas oryzae* pv. oryzicola (Xoc) and *X. oryzae* pv. oryzae (Xoo), respectively. It maps to a 1.09 Mb region of the long arm of chromosome four and segregates as a single dominant locus [24]. Though the molecular mechanism is not yet known, *Xo1* resistance is elicited by any of the several targeted host gene activators, called transcription activator-like (TAL) effectors, injected into the plant cell by Xoc and Xoo; elicitation of resistance does not require the C-terminal TAL effector activation domain, and it is suppressed by N- and C-terminally truncated versions of TAL effectors (truncTALEs; also called iTALEs) found in most Asian strains of the pathogen but missing from examined African strains [24, 28, 29]. The *Xo1* locus overlaps several mapped loci for resistance to BB, including *Xa1*, *Xa2*, *Xa12*, *Xa14*, *Xa17*, *Xa31(t)*, and *Xa38*, that have been isolated from various rice cultivars [30–37]. Of these, *Xa1* has been cloned and encodes an N-terminal, integrated zinc-finger BED [zfBED; 38] domain and uniquely highly conserved, tandem repeats in the LRR region [39]. *Xa1* functions similarly to *Xo1*-mediated resistance, triggered by TAL effectors non-specifically and independent of their ability to activate transcription, and suppressed by truncTALEs [28]. *Xo1* and *Xa1* are together the second discovered example of activation domain-independent TAL effector-triggered resistance, the first being resistance mediated by the tomato Bs4 protein, also an NLR protein, though of the TIR domain type and so far not reported to be suppressed by any truncTALE [40].

Based on the functional similarity of Xo1 to Xa1, and to Bs4, and the fact that the region corresponding to the *Xo1* locus in the rice reference genome (IRGSP-1.0; cv. Nipponbare, which lacks the BLS and BB resistance) [41] contains a complex cluster of seven NLR genes similar to each other (suggesting the potential for rapid evolution), we hypothesized that *Xo1*-mediated resistance in Carolina Gold Select is conferred by an NLR gene at the *Xo1* locus. The Carolina Gold Select genome assembly revealed fourteen such genes at the locus, including a candidate highly similar but not identical to *Xa1*, encoding an N-terminal, integrated zfBED domain and highly conserved, C-terminal, tandem repeats. Herein, in addition to the whole genome assembly, we present a detailed structural and comparative analysis of the *Xo1* candidate and other NLR genes at the *Xo1* locus, and an examination of zfBED-NLR gene content overall across representative species in the tribe Oryzeae.

## RESULTS AND DISCUSSION

### Carolina Gold Select Genome Assembly and Annotation

To generate an assembly made up of large contigs with low error-rate, several assembly methods were used. We found that assembly by Flye [42] using only Nanopore data yielded long contigs but a high consensus error rate. MaSuRCA [43] assembly using both Illumina and Nanopore reads contained more sequence and had a very low consensus error rate, less than 1 error per 10,000 bases. Combining the two assemblies resulted in a reconciled Carolina Gold Select assembly that benefited from both the higher quality consensus sequence and completeness of the MaSuRCA assembly, and the greater contiguity of the Flye assembly. Table 1 lists the quantitative statistics of both assemblies as well as the reconciled assembly. For N50 computations, we used a genome size estimate of 377,689,190 bp, equal to the total size of scaffolds of the final reconciled assembly.

**Table 1.** Quantitative statistics of Carolina Gold Select rice initial assemblies and the final reconciled assembly.

|  |  |  |  |  |  | Consensus<br>error rate<br>(errors per<br>10kb) |
| --- | --- | --- | --- | --- | --- | --- |
| Assembly | N50<br>Contig<sup>a</sup> | N50<br>Scaffold | Output<br>Sequence | # of<br>contigs | # of<br>scaffolds |  |
| **MaSuRCA**<br>(Illumina+Nanopore) | 565,857 | 565,857 | 385,480,701 | 1,942 | 1,942 | <1 |
| **Flye**<br>(Nanopore only) | 1,492,039 | 1,497,653 | 362,619,590 | 649 | 634 | 142 |
| **Reconciled Assembly** | 1,632,109 | 1,719,775 | 377,688,090 | 1,297 | 1,286 | 7 |

<sup>a</sup> Scaffold size of the final assembly (377,689,190 bp) used as genome size for N50 computations.

We found that the Carolina Gold Select assembly mapped to the Nipponbare reference genome with average identity of 98.96%. 350,765,472 bases of the assembly (93%) aligned to 347,609,898 bases (93%) of the reference. The chromosome scaffolding process found 29 breaks in the scaffolds that were apparent mis-assemblies and these were resolved. We call the final chromosomes Carolina_Gold_Select_1.0. The length statistics are provided in Table 2.

**Table 2.** Chromosome sizes for final Carolina Gold Select assembly.

| Chromosome | Base pairs | Number of contigs |
| --- | --- | --- |
| 1 | 43,693,361 | 82 |
| 2 | 33,403,981 | 33 |
| 3 | 36,226,658 | 45 |
| 4 | 26,997,489 | 60 |
| 5 | 32,940,350 | 84 |
| 6 | 29,555,730 | 75 |
| 7 | 32,220,145 | 47 |
| 8 | 27,351,946 | 75 |
| 9 | 22,079,432 | 49 |
| 10 | 26,146,550 | 56 |
| 11 | 29,489,498 | 50 |
| 12 | 25,924,128 | 54 |
| Unplaced | 11,621,710 | 605 |

Protein coding genes were annotated based on the annotation of the reference genome (see Methods). For the 12 chromosomes our mapping process identified and annotated 80,753 gene loci, of which 33,818 have protein coding transcripts. We identified a total of 86,983 transcripts, of which 40,047 are protein coding and have identified CDS features. The total number of bases covered by exons is 52,082,180 bp, or 14.2% of the total length of all 12 chromosomes, whose lengths sum to 366,055,270 bp.

### NLR genes in the Carolina Gold Select Assembly

To identify NLR genes in the Carolina Gold Select genome, we used NLR-Annotator, an expanded version of the NLR-Parser tool [44]. NLR-Annotator does not rely on annotation data and does not mask repetitive regions, facilitating an unbiased analysis of the complete genome including NLR genes [45]. Because the NLR-Annotator pipeline has not been validated in rice, we first ran the pipeline on the well-annotated Nipponbare reference. A total of 518 complete or partial NLR genes were predicted. Genomic locations of these were cross-referenced with a list of 360 Nipponbare NLR genes that were included in a recent analysis [23]; 356 matched. Of the four Nipponbare NLR genes that were not identified by NLR-Annotator, one lacks one or more canonical NLR gene domains based on InterProScan predictions. The other three appear to be complete, however, indicating an overall NLR-Annotator detection success rate of 99.2% (Additional file 1). NLR-Annotator identified some complete NLR genes in the Nipponbare genome distinct from the 356; these may represent previously undetected NLR genes, pseudogenes, or false positives.

Running the Carolina Gold Select assembly through the NLR-Annotator pipeline identified 529 complete or partial NLR genes. The Carolina Gold Select NLR genes are organized similarly to those of Nipponbare, occurring irregularly across the 12 chromosomes, with a large proportion occurring on chromosome 11 (Fig. 1 and Additional file 2). This similarity in number and genomic distribution of NLR genes provides support for the integrity of the Carolina Gold Select genome assembly.

**Figure 1.**
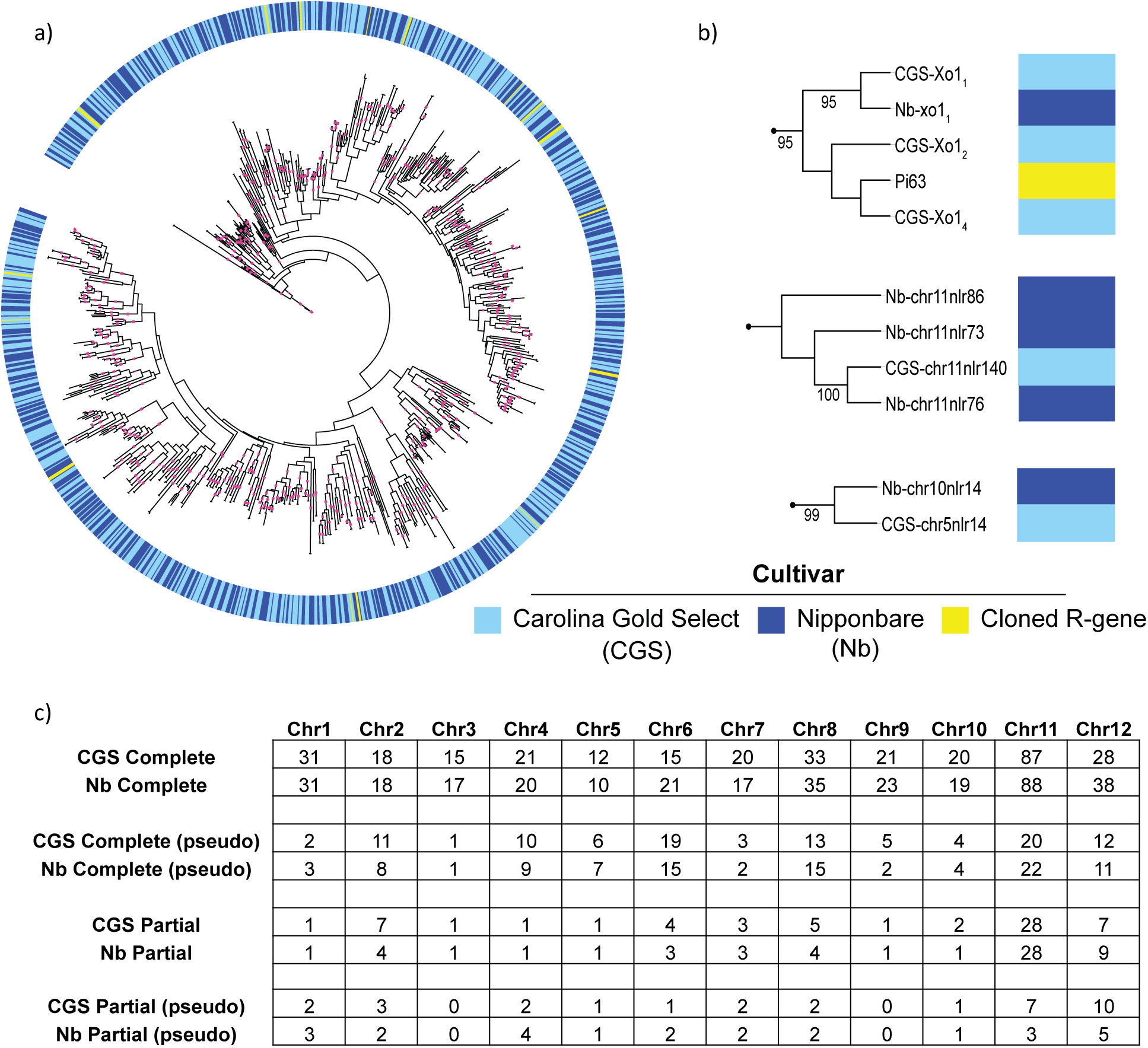
NLR proteins encoded in Carolina Gold Select in relation to Nipponbare and selected *R* genes. (a) Maximum likelihood tree of encoded NB-ARC domains of NLR genes in Carolina Gold Select (436) and Nipponbare (427), as predicted by NLR-Annotator. Incomplete NLR genes are not included in the phylogeny. Fifteen cloned resistance genes are included for reference. Branches with bootstrap support greater than 80 are indicated with pink squares. Interactive tree available at http://itol.embl.de/shared/acr242. NB-ARC domain sequences available in Additional file 3. (b) Examples of expansion (top), contraction (middle) and transposition (bottom) of NLR genes in Carolina Gold Select relative to Nipponbare; bootstrap values greater than 80 are displayed. Further details available in Additional file 4. (c) Number and chromosomal distribution of all NLR-Annotator predicted NLR genes in Carolina Gold Select and Nipponbare.

To determine relationships between and among Nipponbare and Carolina Gold Select NLR genes, amino acid sequences of the central NB-ARC domain for all complete NLR genes were used to generate a maximum likelihood phylogenetic tree. NB-ARC domains from 15 cloned, NLR-type, rice resistance genes (Additional file 3) were included to identify potential orthologs in Carolina Gold Select. Although the total number of predicted NLR genes is similar between the two cultivars, the resulting tree revealed 32 expansions and 34 contractions within NLR gene clusters in Carolina Gold Select relative to Nipponbare, as well as 3 transpositions and 4 transpositions combined with expansion or contraction (Fig. 1 and Additional file 4). Seven of the cloned resistance genes (*Pib*, *Pik2*, *Pi63*, *Pi2*, *RGA5*, *Pi36*, and *Pi37*) cluster with expanded or contracted NLR gene groups. The observed differences in NLR gene content in the two closely related cultivars is consistent with previous comparative analyses demonstrating that NLR gene families evolve rapidly and are characterized by presence-absence variation [18–20].

### Expansion at the Carolina Gold Select *Xo1* locus

We next examined the *Xo1* locus. We extracted the region of the Carolina Gold Select assembly that corresponds to the 1.09 Mb Nipponbare *Xo1* mapping interval [24] and found that it spans a much larger region, 1.30 Mb, that includes a 182 kb insertion (Fig. 2). It is unclear if this relative expansion is unique to a particular subgroup of *O. sativa* cultivars, but it is not present in the long-read (PacBio) assembly of *O. sativa* indica cultivar IR8 (Additional file 5) [46]. Hereafter, we refer to the region in Nipponbare, which as noted lacks the resistance to BLS and BB, as *Nb-xo1* and to the region in Carolina Gold Select as *CGS-Xo1*.

**Figure 2.**
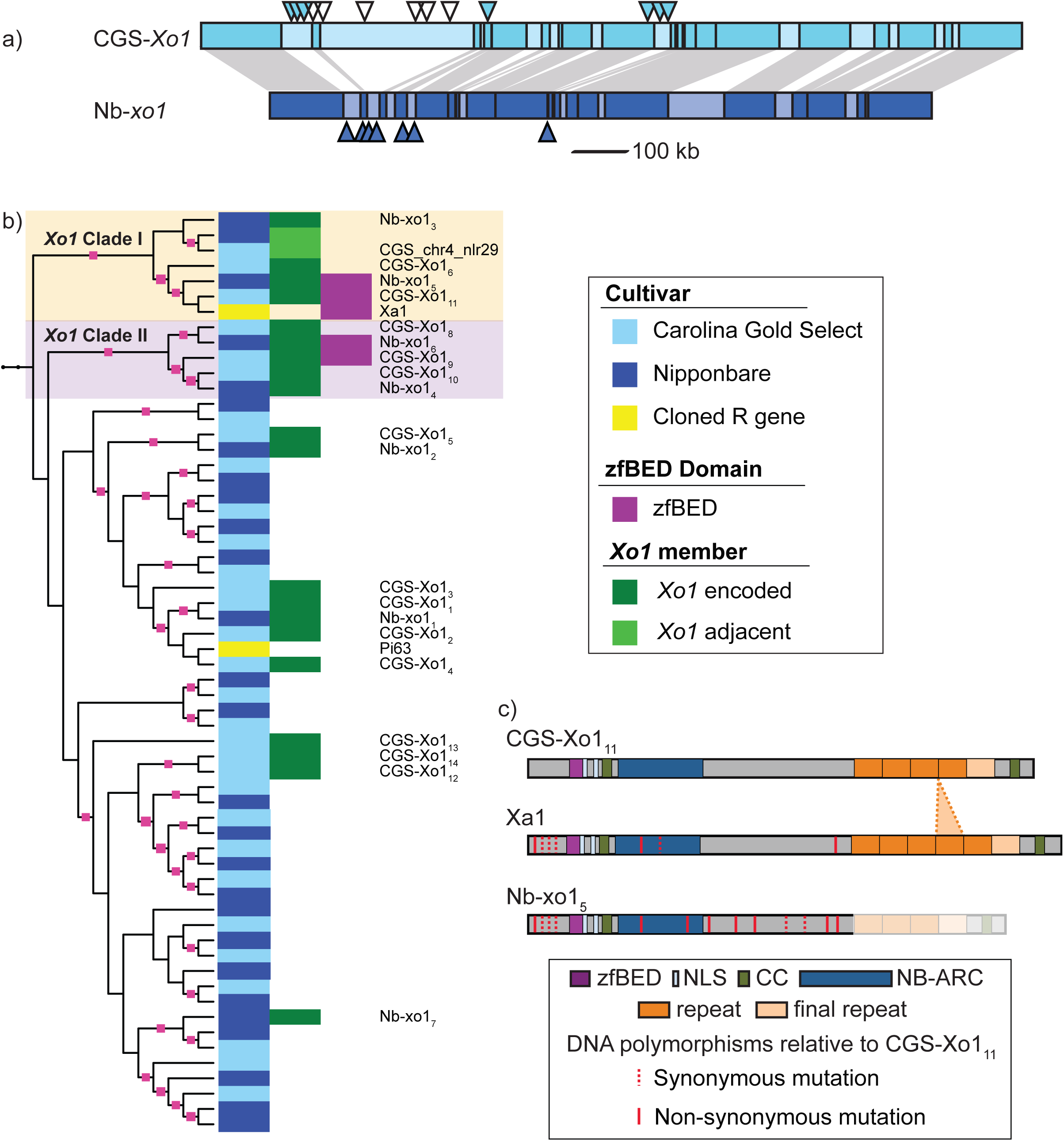
Expansion at the Carolina Gold Select *Xo1* locus and identification of an ***Xo1* candidate**. (a) Comparison of the *Xo1* locus in Carolina Gold Select and in Nipponbare. Triangles indicate positions of NLR genes predicted by NLR-Annotator, designated from left to right as *CGS-Xo1_1_*through *CGS-Xo1_14_* in Carolina Gold Select and *Nb-xo1_1_*through *Nb-xo1_7_* in Nipponbare. Filled triangles indicate NLR genes expressed in leaf tissue during infection (see text and Additional file 14). (b) An excerpt of the phylogenetic tree from Figure 1a containing the NLR genes at the *Xo1* locus and two known resistance genes, *Xa1* and *Pi63*. NLR genes encoding an integrated zfBED domain fall into two clades, which we designate as *Xo1* clades I and II. Branches with bootstrap support greater than 80 are indicated with pink squares. Interactive tree available at http://itol.embl.de/shared/acr242. (c) Cartoon alignment of predicted products of *CGS-Xo1_11_*, *Xa1*, and *Nb-xo1_5_*showing the zfBED domains, nuclear localization signals (NLS), coiled coil domains (CC), NB-ARC domains, tandem repeats, and final repeats. Synonymous and nonsynonymous nucleotide substitutions in relation to CGS-Xo1_11_ are indicated by dashed and solid red lines respectively. Further comparisons can be found in Additional file 7.

We mapped the NLR-Annotator output for Carolina Gold Select and Nipponbare onto the locus (Fig. 2). There are 14 predicted NLR genes at *CGS-Xo1*, which we name *CGS-Xo1_1_*through *CGS-Xo1_14_*. There are seven at *Nb-xo1*, matching the annotation of the reference genome; we refer to these as *Nb-xo1_1_*through *Nb-xo1_7_* (Additional file 1). The NLR genes are not evenly distributed across the locus, but instead occur in clusters, consistent with the previous observation that only 24.1% of rice NLR genes occur as singletons [15].

### Identification of an *Xo1* candidate

Having delineated NLR gene content at the *Xo1* locus, we then sought to identify a candidate or candidates for the *Xo1* gene itself. First, using RNA sequencing (RNAseq), we asked which of the 14 predicted *CGS-Xo1* NLR genes are expressed in rice leaves following inoculation with an African strain of Xoc, that strain expressing a truncTALE, or a mock inoculum. The data provided evidence for expression of 8 of the 14 NLR genes (Fig. 2). In contrast, each of the NLR genes at the locus in Nipponbare is expressed, based on previously obtained RNAseq data from leaves inoculated with the same African strain of Xoc [47]. The lack of expression data for nearly half the NLR genes at the CGS-Xo1 locus led us to question whether the observed expansion at *CGS-Xo1* is an artifact of the assembly. To determine whether this is the case, we mapped all Nanopore reads to the assembly using BLASR [48], picked one best alignment for each read, and then examined the read coverage in the vicinity of the *CGS-Xo1* locus. The Nanopore reads covered the region with average depth of 21x, varying from 18x to 25x, providing robust support for the assembly. Thus, we considered the eight NLR genes expressed under the tested conditions to be candidates for *Xo1*; the other six may be non-functional, or expressed under different conditions or tissues. We cannot rule out the possibility that the resistance is conferred by one or more of the non-NLR genes at the locus, but none of the annotations for those genes suggests a role in immunity (Additional file 6).

Next, we inspected the NB-ARC domain-based phylogenetic tree and observed that the susceptible cultivar Nipponbare and the resistant cultivar Carolina Gold Select have one NLR gene each, *Nb-xo1_5_*and *CGS-Xo1_11_*, that group closely with *Xa1*, the cloned BB resistance gene functionally similar to *Xo1* (Fig.1c). Several additional NLR proteins encoded at the *Nb-xo1* and *CGS-Xo1* loci fall into the same or a closely related clade. We call these *Xo1* clade I and *Xo1* clade II, respectively. They both reside in major integration clade (MIC) 3 defined by Bailey *et al.* [23]. Using the *Xa1* coding sequence as a guide, we extracted and aligned the corresponding sequences from *Nb-xo1_5_* and *CGS-Xo1_11_*(Fig. 2c). The MSU7 [41] gene model for *Nb-xo1_5_* (LOC_Os04g53120) indicates that there is an intron downstream of the repeats, however, the sequence in the predicted intron aligns well to *CGS-Xo1_11_* and *Xa1* coding sequence and therefore seems likely to be a mis-annotation. Thus, in our alignment we included it as coding sequence. Based on the Carolina Gold Select and Nipponbare genomic sequences, each of the coding sequences corresponds to three exons. The first is 307 bp and encodes no detectable, known protein domains. The second, 310 bp, encodes a non-canonical, integrated, 49 amino acid (aa) zfBED domain and a predicted, 9 aa nuclear localization signal (NLS). The third exon, the longest, encodes a second predicted 9 aa NLS, a 21 aa coiled coil (CC) domain, a 288 aa NB-ARC domain, the LRR region, and a second, C-terminal, 21 aa coiled coil domain. There are very few differences in the three genes upstream of the LRR-encoding region. In fact the zfBED domain, 2 NLSs, and first coiled coil domain are 100% conserved at the nucleotide level. There is a single amino acid difference between the *CGS-Xo1_11_* and *Xa1* NB-ARC domains, and two, distinct differences in that domain between *CGS-Xo1_11_*and *Nb-xo1_5_*. In *Nb-xo1_5_*, the MHD triad, which has a role in NLR activation [49], has a M to V substitution. This substitution seems unlikely to be functionally relevant, however, as VHD has been observed in several functional CC-NLR proteins [50].

The LRR regions of *CGS-Xo1_11_* and *Nb-xo1_5_* share with *Xa1* the striking feature of highly conserved, tandem repeats in the LRR region. Though LRR regions are partially defined by their repetitive aa sequence, typically the repeats are polymorphic. The repeats within the *CGS-Xo1_11_*, *Nb-xo1_5_*, and *Xa1* LRR regions, each 93 aa (279 bp) in length, are nearly identical to one another. To explore this feature further, we analyzed all predicted NLR genes from the Nipponbare reference and the Carolina Gold Select assembly and found that, among the >1000 sequences, nearly identical LRRs are found only in NLR proteins encoded at the *Nb-xo1*/*CGS-Xo1* locus, though not all NLR genes at these loci encode such repeats. *Xa1*, *CGS-Xo1_11_*, and *Nb-xo1_5_*, despite sharing the feature, differ in the number and conservation of their repeats. *Xa1* has five full repeats while *CGS-Xo1_11_* has four and *Nb-xo1_5_* three (Fig. 2c and Additional file 7). Each gene encodes an additional, less conserved, final repeat. Intra- and inter-repeat comparison shows that *CGS-Xo1_11_*and *Xa1* align well while *Nb-xo1_5_* is more divergent (Additional file 7). Overall, the sequence relationships suggest that *CGS-Xo1_11_*is the *Xo1* gene. Functional analysis will be required to test this prediction definitively.

### *CGS-Xo1_11_*-like genes encoded in Oryzeae

The differences we observed in the presence of the zfBED domain and of the nearly identical repeats among NLR proteins encoded at the CGS-*Xo1* and Nb-*xo1* loci prompted us to characterize diversity of these features across the Oryzeae tribe. We ran the NLR-Annotator pipeline on the genomes of *Leersia perrieri, O. barthii, O. glaberrima, O. glumaepatula, O. brachyantha, O. meridinalis, O. nivara, O. punctata, O. rufipogon, O. sativa* IR8, and *O. sativa* Aus N22 [46, 51, 52]. All NB-ARC domains identified were added to those of Nipponbare and Carolina Gold Select. These >5,000 sequences were used to generate an Oryzeae NLR gene maximum likelihood phylogenetic tree (Additional file 8). Two distinct sister clades in the tree respectively include the previously identified Carolina Gold Select and Nipponbare *Xo1* clade I and II NLR genes. Full sequences of the NLR genes represented in these expanded *Xo1* clades I and II were extracted and examined for the presence of a zfBED domain, additional IDs, and nearly identical repeats (Fig. 3 and Additional file 9). NLR genes from each genome are found in each clade; however, not all clade I and II NLR genes encode a zfBED domain, and no NLR genes of *O. brachyantha* do. Nearly identical repeats are found only in NLR genes with a zfBED domain, though there are several zfBED-NLR genes without them. A zfRVT domain (zinc-binding region of a putative reverse transcriptase; Pfam 13966) was predicted in four *Xo1* clade I Oryzeae NLR genes as well as one of the wheat Yr alleles from *Xo1* clade II. The zfRVT domain has been detected in previous NLR gene surveys [22]. Most of the NLR genes in the two clades reside in the *Xo1* locus on chromosome four, however there are six, all from *Xo1* clade II, that are on other chromosomes; this is consistent with research demonstrating that transposition events are common during evolution of NLR gene families [53].

**Figure 3.**
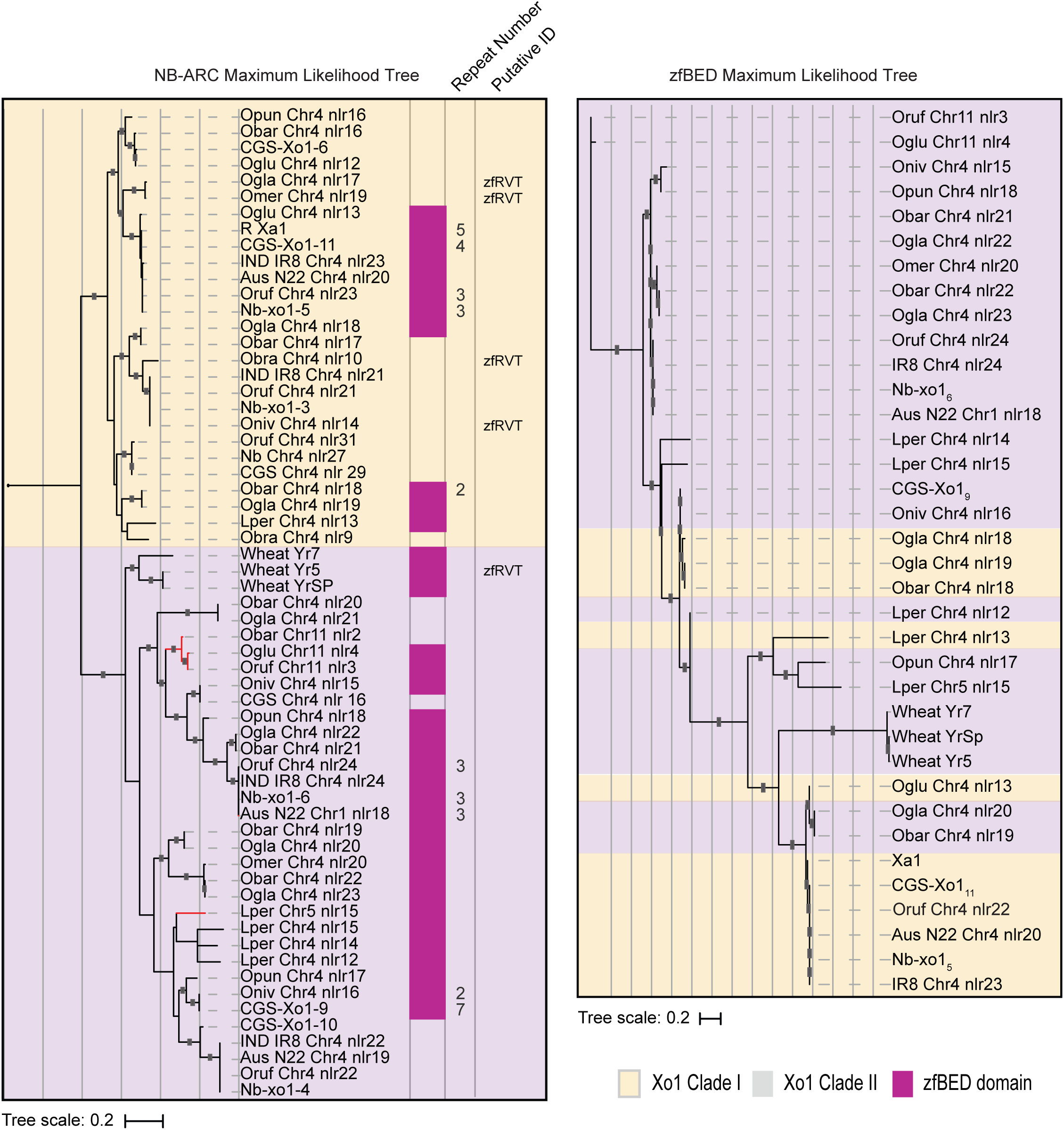
zfBED-NLR proteins across the Oryzeae. (a) *Xo1* clade I and II from an NB-ARC domain-based maximum likelihood tree of 5104 predicted NLR proteins from representative Oryzeae genomes. Numbers of tandem 279 bp C-terminal repeats, where present, are given. Additional detected, non-canonical NLR gene motifs are noted. Red branches correspond to NLR genes not on chromosome four. Full Oryzeae tree in Additional file 8 and interactive tree available at http://itol.embl.de/shared/acr242. (b) Maximum likelihood tree of the 36 predicted Oryzeae zfBED-NLR proteins based on the zfBED domain amino acid sequence (zfBED sequences and nucleotide tree in Additional files 10 and 11). Branches with bootstrap support greater than 80 are indicated with grey squares. Interactive trees available at http://itol.embl.de/shared/acr242.

The presence of closely related zfBED-NLR genes across diverse Oryzeae species suggests that the integration of the zfBED domain preceded Oryzeae radiation. This inference is consistent with a recent analysis that identified NLR genes encoding N-terminal zfBED domains in several monocot species including *Setaria italica*, *Brachypodium distachyon*, *Oryza sativa*, *Hordeum vulgare*, *Aegilops tauschii*, *Triticum urartu*, and *Triticum aestivum*, though no zfBED-NLR genes were detected in *Sorghum bicolor* or *Zea mays* [23]. ZfBED-NLR genes have also been detected in dicots, with as many as 32 reported in poplar (*Populus trichocarpa*) [54]. A more recent analysis that includes *P. trichocarpa* detected 26 zfBED-NLR genes, of which 24 have the same architecture as *CGS-Xo1_11_*, with the zfBED domain encoded upstream of the NB-ARC and LRR domains [22]. Nevertheless, it is unclear if all zfBED-NLR genes arose from a single integration, or if the integration has occurred independently in the monocot and dicot lineages. Distribution among dicots seems limited, and a recent delineation of the *Arabidopsis* pan ‘NLR-ome’ generated from 65 accessions found none [55].

Three alleles of a zfBED-NLR gene in wheat, *Yr5*, *Yr7*, and *YrSP*, were recently shown to provide resistance to different strains of the stripe rust pathogen, *Puccinia striiformis* f. sp. tritici [56]. The *Yr5*/*Yr7*/*YrSP* syntenic region in the Nipponbare genome, determined by the authors of that study, overlaps *Nb-Xo1_5_*. When added to the Oryzeae tree, the NB-ARC domains of the wheat rust resistance alleles cluster with *Xo1* clade II (Fig. 3). It is remarkable that these evolutionarily-related NLR genes with similar non-canonical N-terminal fusions provide resistance to two pathogens from different kingdoms of life. In this context it is also worth noting that the poplar *MER* locus for *Melampsora larica-populina* rust resistance was reported to contain 20 of the 32 poplar zfBED-NLR genes [54].

It has been demonstrated in rice and *Arabidopsis* that IDs in NLR proteins can act as decoys for pathogen effector proteins such that their interaction with an effector activates the NLR protein and downstream defense signaling [22, 57–59]. If this were the case for the zfBED domain, we might expect to see distinct signatures of evolution in the zfBED and NB-ARC domains. We extracted the zfBED domains from 33 Oryzeae zfBED-NLR genes as well as the three wheat *Yr* alleles and created a tree to determine if they would cluster into two sub-groups, similarly to the NB-ARC domains. They do not, even when the tree is generated from the nucleotide sequences (Fig. 3 and Additional files 10 and 11). The zfBED domain of *Xo1* clade I and II NLR genes thus appears to be under distinct selective pressures from the NB-ARC domain. Alternatively, the discordance between the NB-ARC and zfBED trees may be evidence of domain swapping, as has been reported for other integrated domain-encoding NLR genes [60].

The role or roles of the zfBED domain remain unclear. The observations that *Yr7*, *Yr5*, and *YrSP* have identical zfBED sequences but recognize different pathogen races [56] and that *Xa1*, *CGS-Xo1_11_*, and *Nb-xo1*_5_ encode identical zfBEDs, does not support the model of this domain being a specificity-determining decoy. Rather, it may have a role in downstream signaling, a role in localization, or some other role. Mechanisms might include dimerization, recruitment of other interacting proteins, or DNA binding.

### The Carolina Gold Select *Xo1* locus contains a rice blast resistance gene

In the NB-ARC domain-based tree (Fig. 1), *CGS-Xo1_2_* and *CGS-Xo1_4_* group with rice blast resistance gene *Pi63*, originally cloned from rice cultivar Kahei [61, 62]. Direct sequence comparison revealed that *Xo1_4_*, which is expressed (Fig. 2a), is *Pi63*: the genomic sequences, including 3 kb upstream of the gene bodies, are 100% identical (not shown). Modern US rice varieties, some of which descend from initial Carolina Gold populations, contain several blast resistance genes including *Pik-h*, *Pik-s*, *Pi-ta*, *Pib*, *Pid*, and *Pi2* [reviewed in 63], but each of these genes was introduced into the US germplasm from Asian cultivars, and none resides on chromosome four. Our discovery of *Pi63* in Carolina Gold Select reveals that this variety may be a useful genetic resource for further strengthening US rice blast resistance.

The presence of blast and blight resistance at the *Xo1* locus in Carolina Gold Select is reminiscent of *O. sativa* japonica cultivar Asominori. Asominori is the source of the blast resistance gene *PiAs(t)* and the BB resistance gene *Xa17*, and both of these genes, though not yet cloned, map to the *Xo1* region of chromosome four. *Xa17*, previously *Xa1-As(t)*, has a similar resistance profile to *Xa1* but provides resistance at both seedling and adult stages; *Xa1* is unstable at the seedling stage [33]. *PiAs(t)* and *Xa17* are closely linked to a polyphenol oxidase (PPO) gene, the activity of which can be detected by treating seeds with phenol [33]. This seed-treatment assay has been used as a surrogate to track the blight and blast resistance genes during crosses [32, 33]. In the Carolina Gold Select genome assembly, *CGS-Xo1_4_* (*Pi63*) and *CGS-Xo1_11_* are separated by 270 kb, and a PPO gene resides an additional 175 kb downstream. However, the Carolina Gold Select PPO gene sequence has a 29 bp loss-of-function deletion common in japonica cultivars [64]. The seed treatment assay confirmed absence of PPO activity (Additional file 12). It seems likely that the genomic arrangement at the Asominori blight and blast resistance locus is similar to that in Carolina Gold Select, though with an intact PPO gene. Our results illustrate that while the seed treatment assay may be useful to track resistance at the *Xo1* locus in some cases, such as crosses with Asominori, in others it may not, due to a loss of function mutation in the linked PPO gene. More broadly, our results demonstrate the ability to make phenotypic predictions based on the Carolina Gold Select assembly.

## CONCLUSIONS

In this study, whole genome sequencing using Nanopore long reads along with Illumina short reads delineated a complex, NLR gene-rich region of interest, the *Xo1* locus for resistance to BLS and BB, in the American heirloom rice variety Carolina Gold Select. This revealed an expansion at the locus relative to the reference (Nipponbare) genome and allowed identification of an *Xo1* gene candidate based on sequence similarity to the functionally similar, cloned *Xa1* gene, including an intergrated zfBED domain and nearly identical repeats. Analysis of NLR gene content genome-wide and comparisons across representative members of the Oryzeae and other plant species identified two sub-clades of such NLR genes, varying in the presence of the zfBED domain and the number of repeats. The results supported the conclusion of Bailey *et al.* [23] that the zfBED domain was integrated prior to the differentiation of the Oryzeae, possibly before divergence of monocots and eudicots, and revealed that the zfBED domain has been under different selection from the NB-ARC domain. The results also provided further evidence that the zfBED domain can be identical not only among resistance alleles with different pathogen race specificities but also between resistance genes that recognize completely different pathogens [56]. Considering *CGS-Xo1_11_* and *Nb-Xo1_4_*, the results also suggested that the zfBED domain can be identical between functional and non-functional, expressed resistance gene alleles. Finally, the genome sequence uncovered a known rice blast resistance gene at the *Xo1* locus and a loss of function mutation in a linked, PPO gene. The latter breaks the association of PPO activity with BB and blast resistance that has been the basis of a simple, seed staining assay for breeders to track the resistance genes in some crosses.

Our study illustrates the feasibility and benefits of high quality, whole genome sequencing using long- and short-read data to resolve and characterize individual, complex loci of interest. It can be done by small research groups at relatively low cost: our sequencing of the Carolina Gold Select genome used data generated from a single Illumina HiSeq2500 lane and two ONT MinION flowcells. Because long-read sequencing technologies and base-calling continue to improve, it seems likely that high quality assemblies from long-read data alone will become routine. The long-read data enabled us to identify and characterize the expansion of NLR genes at the *Xo1* locus.

Such presence/absence variation across genotypes is hard if not impossible to determine definitively by only short-read sequencing. The long-read data, with short-read error correction, also allowed us to define the number and sequences of nearly identical repeats in the *Xo1* gene candidate *CGS-Xo1_11_* and genes like it in Carolina Gold Select. Indeed, we caution that, in short-read assemblies, sequences of *CGS-Xo1_11_*homologs and other such repeat-rich genes, or repeat-rich intergenic sequences, should be interpreted with care, due to the possibility of artificially collapsed, expanded, or chimeric repeat regions.

Cataloging NLR gene diversity in plants is of interest for resistance gene discovery, for insight into NLR gene evolution, and for clues regarding the functions of IDs. Sequence capture by hybridization approaches, such as RenSeq, have been developed and applied to catalog NLR genes in representative varieties of several plant species [65–71], but these depend on *a priori* knowledge to design the capture probes and thus may miss structural variants. Also, they do not reveal genomic location, recent duplications, or arrangement of the genes, information necessary to investigate evolutionary patterns. Sequence capture of course also misses integrated domains or homologs encoded in non-NLR genes, precluding broader structure-function and evolutionary analyses. Sequence capture is nevertheless likely to continue to play an important role in organisms with large, polyploid, or otherwise challenging genomes.

The Carolina Gold Select genome sequence is among a still relatively small number of high quality assemblies for rice and the first of a tropical japonica variety. The identification of an *Xo1* candidate is a significant step toward cloning and functional characterization of this important gene and will facilitate investigation of the role(s) of the integrated zfBED domain in NLR gene-mediated resistance. The Carolina Gold Select genome assembly will be an enabling resource for geneticists and breeders to identify, characterize, and make use of genetic determinants of other traits of interest in this historically important rice variety.

## METHODS

### Genomic DNA Extraction and Nanopore Sequencing

Carolina Gold Select seedlings were grown in LC-1 soil mixture (Sungro) for three weeks in PGC15 growth chambers (Percival Scientific) in flooded trays with 12-hour, 28°C days and 12-hour, 25°C nights. Three weeks after planting leaf tissue was collected and snap frozen in liquid nitrogen.

Genomic DNA was extracted from 250 mg of frozen leaf tissue with the QIAGEN g20 column kit with 0.5 mg/ml cellulase included in the lysis buffer. Eluted DNA was cleaned up with 1 volume of AMPure XP beads (Beckman-Coulter). To attain the recommended ratio of molar DNA ends in the Nanopore library prep the genomic DNA was sheared with a Covaris g-TUBE for one minute at 3800 RCF on the Eppendorf 5415D centrifuge. A 0.7x volume of AMPure XP beads was used for a second clean-up step to remove small DNA fragments. Sheared DNA was analyzed on a NanoDrop spectrophotometer (Thermo Fisher) to determine A260/280 and A260/230 ratios, and quantified using the Qubit dsDNA BR (Broad Range) assay kit (ThermoFisher).

Fragment length distribution was visualized with the AATI Fragment Analyzer (Agilent). Sheared genomic DNA was used as input into the Nanopore LSK108 1D-ligation library prep kit, then loaded and run on two R9.4.1 MinION flow cells. Raw reads for both flow cells were base-called with Albacore v2.3.0. Amounts of DNA at each step of the workflow can be found in Additional file 13. Run metrics were calculated using scripts available at https://github.com/roblanf/minion_qc.

### Illumina Sequencing

Genomic DNA was isolated from leaf tissue of a single Carolina Gold Select plant using the Qiagen DNEasy kit. Libraries were prepared as described [72], using the Illumina TruSeq kit with an insert size of ∼380 bp. Two × 100-bp paired-end sequencing was carried out on an Illumina HiSeq 2500.

### Sequence Assembly

Reads were assembled using default settings with two different assembly programs, MaSuRCA version 3.2.7 [43] and Flye version 2.4.1[42], followed by reconciliation of the results to produce an initial contig/scaffold assembly of the genome, CG_RICE_0.9. In reconciliation we followed the procedure described in [73]. We merged the contigs from the more contiguous Flye assembly with MaSuRCA contigs by mapping the assemblies to each other using Mummer4 [74], then filtering the alignments for reciprocal best hits and looking for alignments longer than 5000 bp where one assembly merged the contigs of the other. This resulted in longer merged contigs with the relatively low-quality consensus of the Flye assembly. We then aligned the high quality MaSuRCA assembly contigs to the merged contigs using Mummer4, filtered for unique best alignments for each contig, and replaced the consensus of the merged contigs with MaSuRCA consensus, resulting in a highly contiguous, merged assembly with low consensus error rate. Consensus error rate was computed using the script ‘evaluate_consensus_error_rate.sh’ distributed with MaSuRCA, which was created following [75]; this script maps the Illumina data to the assembly using bwa [76], and then calls short sequence variants using freebayes software [77]. A sequence variant at a site in a contig sequence is an error in consensus if all Illumina reads disagree with consensus at the site, and there are at least three Illumina reads that agree on an alternative. Sequence variants are SNPs and short insertions/deletions. Total number of errors is the total number of bases in error variant calls, and the error rate is computed as total number of errors divided by the sequence size.

Following the completion of the assembly, we used the Nipponbare rice reference genome IRGSP-1.0 (NCBI accession GCF_001433935) [41] to order and orient the assembled scaffolds on the chromosomes using the MaSuRCA chromosome scaffolder tool, publicly available as part of the MaSuRCA distribution starting with version 3.2.7.

### Reference-based Annotation

We annotated the 12 assembled chromosome sequences by aligning the transcripts from the rice annotation produced by the International Rice Genome Sequencing Project (IRGSP) and the Rice Annotation Project Database (RAP-DB) [41] We used release 1.0.40 of the annotation file for *Oryza sativa* made available by Ensembl Plants [78]. We aligned the DNA sequences of these transcripts to our assembled chromosomes using GMAP [79]. The resulting exon-intron mappings were further refined for transcripts annotated as protein coding, as follows. For each protein-coding transcript in our assembled chromosomes, we extracted the transcript sequence using gffread (http://ccb.jhu.edu/software/stringtie/gff.shtml) and aligned it with the protein sequence from the IRGSP annotation to identify the correct start and stop codon locations. These protein-to-transcript sequence alignments were performed using blat [80], followed by a custom script that projected the local CDS coordinates back to the exon mappings on our assembled chromosome sequences, to complete the annotation of the protein-coding transcripts.

### RNA Extraction and Sequencing

Three-week old Carolina Gold Select seedlings grown under the conditions described above were syringe-infiltrated with an OD_600_ 0.4 suspension of African Xoc strain CFBP7331 carrying a plasmid-borne copy of the truncTALE gene *tal2h* or empty vector [29], or mock inoculum (10 mM MgCl_2_). Each leaf was infiltrated at 20 contiguous spots starting at the leaf tip. Inoculated tissue was harvested 24-hours post-infiltration, before the hypersensitive reaction manifested for CFBP7331 with empty vector. The experiment was repeated three times. RNA was extracted from the replicates with the QIAeasy RNA extraction kit (Qiagen) and submitted to Novogene Biotech for standard, paired-end Illumina sequencing. For Nipponbare, previously generated RNAseq data was used (Accessions SRX978730, SRX978731, SRX978732, SRX978723, SRX978722, and SRX978721, Short Read Archive of the National Center for Biotechnology Information). These data were generated from leaf tissue collected 48 hours after inoculation with CFBP7331 [47].

### NLR gene expression analysis

Genomic sequences of all NLR-Annotator-identified genes plus 1 kb upstream and 1kb downstream were extracted and used to generate indices for Nipponbare and Carolina Gold Select. The additional sequences on each end were included in an effort to capture the entire transcript while avoiding transcripts for any genes encoded nearby. To quantify expression, we used the ‘quant’ function in Salmon [81], mapping reads to the appropriate index. *Xo1* NLR genes with >500 Transcripts per Kilobase Million were considered expressed (Additional file 14).

### NLR Gene Identification and Phylogenetic Analysis

NLR gene signatures were detected with NLR-Annotator [82] using a sequence fragment length of 20 kb with 5 kb overlaps. NLR-Annotator predictions for Nipponbare were compared to previously annotated NLR genes using BED-tools intercept [83].

Encoded NB-ARC domains for all Nipponbare and Carolina Gold Select NLR genes and for the additional rice NLR genes were extracted using NLR-Annotator and aligned using Clustal-omega [84] with default settings. Maximum likelihood trees were generated with RAxML v8.2.12 [85] with 100 bootstraps and visualized using the Interactive Tree of Life (iTOL) tool [86]. This pipeline was repeated for the representative group of Oryzeae genomes.

Integrated domains outside of the canonical NLR gene structure were detected by running the NLR-Annotator-identified genes plus the 5 kb 5′ and 5 kb 3′ flanking sequences (Additional file 15) through Conserved Domain BLAST [87] using default parameters. Domains >2 kb from a known NLR domain were considered likely false positives and disregarded. Domains deemed likely to be annotations of LRR sub-types were also excluded.

### Tandem Repeat Characterization

Self-comparison dotplots were used to determine whether NLR-Annotator-identified genes in Nipponbare and Carolina Gold Select contain nearly identical repeats. In order to define repeat units in a standardized way, *Xo1* clade I and II NLR gene sequences were extracted and submitted to Tandem repeats finder with default parameters [88]. WebLogos for aligned repeats of *CGS-Xo1_11_*, *Xa1*, and *Nb-xo1_6_* were generated using WebLogo3 [89].

## Supporting information

Additional_file_14_Xo1_TPM

Additional_file_13_NanoporeMetrics

Additional_file_12_PPO_figure

Additional_file_11_zfBED_nt_tree

Additional_file_10_BEDsequences

Additional_file_9_Xo1_CDBLAST

Additional_file_8_Maximum_likelihood_tree_Oryzeae

Additional_file_7_TandemRptAlignment

Additional_file_6_Annotated_Genes_Xo1

Additional_file_5_IR8vsCGS_dotplot

Additional_file_4_ExpContrTrans

Additional_file_3_AllNBARC

Additional_file_2_CGSNLRAnnotator

Additional_file_1_Nb_NLRAnnotator

## Acknowledgements

The authors thank M. Hutin and current members of the Bogdanove laboratory for helpful discussion. The authors also gratefully acknowledge contributors to Protocols.io, which was useful in optimizing DNA extraction and library preparation for the Nanopore sequencing.

## Availability of data and materials

Carolina Gold Select germplasm: USDA:GRIN database ID GSOR301024 BioSample SAMN10380581 BioProject PRJNA503892 Nanopore and Illumina genomic reads: pending at time of submission of this manuscript Illumina RNAseq reads: SRX6087556, SRR9320041, SRX6087557, SRR9320040, SRX6087558, SRR9320039, SRX6087559, SRR9320038, SRX6087560, SRR9320037, SRX6087561, SRR9320036, SRX6087562, SRR9320035, SRX6087563, SRR9320034, SRX6087564, SRR9320033

## Authors’ contributions

AR, LT, and AB conceived the study. AR carried out nanopore sequencing. RM and LT carried out Illumina sequencing, with contributions from MP and JL. AZ, GP, and SS assembled and annotated the genome. AR and MM identified NLR genes and conducted the RNAseq analysis. AR, MM, and AB analyzed phylogenetic data. AR drafted the manuscript and all authors contributed to the final version.

## ADDITIONAL FILES

Additional file 1 – NLR-Annotator output for Nipponbare cross-referenced with the MSU 7 annotation

Additional file 2 – NLR-Annotator output for Carolina Gold Select

Additional file 3 – NB-ARC domain sequences used to generate the phylogenetic tree in Fig. 1

Additional file 4 – Expansion, contraction, and transposition of NLR gene clusters in Carolina Gold Select relative to Nipponbare

Additional file 5 – Dotplot comparison of the *Xo1* locus in Carolina Gold Select and IR8

Additional file 6 – Annotated genes at the Carolina Gold Select *Xo1* locus Additional file 7 – Tandem repeats in *CGS-Xo1_11_*, *Xa1*, and *Nb-xo1_5_* Additional file 8 – Maximum likelihood tree of NLR genes across Oryzeae. Additional file 9 – Integrated domains detected with CD BLAST

Additional file 10 - zfBED domain sequences from the zfBED-NLR genes across the Oryzeae represented in Fig. 3.

Additional file 11 – Maximum likelihood tree of *Xo1* zfBED nucleotide sequences Additional file 12 – Polymorphism at the polyphenol oxidase gene linked to BB and blast resistance genes at the *Xo1* locus

Additional file 13: Nanopore DNA sequencing metrics

Additional file 14: TPM values for Nipponbare and Carolina Gold Select NLR-Annotator predicted genes

Additional file 15: NLR gene sequences plus 5 kb on either side including integrated domains

## REFERENCES CITED

1. Sedlazeck FJ, Lee H, Darby CA, Schatz MC: Piercing the dark matter: bioinformatics of long-range sequencing and mapping. Nat Rev Genet 2018, 19:329–346.

2. Payne A, Holmes N, Rakyan V, Loose M: BulkVis: a graphical viewer for Oxford nanopore bulk FAST5 files. Bioinformatics 2018:bty841.

3. Michael TP, Jupe F, Bemm F, Motley ST, Sandoval JP, Lanz C, Loudet O, Weigel D, Ecker JR: High contiguity Arabidopsis thaliana genome assembly with a single nanopore flow cell. Nat Comm 2018, 9:541.

4. Schmidt MH, Vogel A, Denton AK, Istace B, Wormit A, van de Geest H, Bolger ME, Alseekh S, Mass J, Pfaff C, et al: De novo assembly of a new Solanum pennellii accession using nanopore sequencing. Plant Cell 2017, 29:2336–2348.

5. Jupe F, Rivkin AC, Michael TP, Zander M, Motley ST, Sandoval JP, Slotkin RK, Chen H, Castanon R, Nery JR, Ecker JR: The complex architecture and epigenomic impact of plant T-DNA insertions. PLoS Genet 2019, 15:e1007819.

6. Giolai M, Paajanen P, Verweij W, Witek K, Jones JDG, Clark MD: Comparative analysis of targeted long read sequencing approaches for characterization of a plant’s immune receptor repertoire. BMC Genomics 2017, 18:564.

7. Ossowski S, Schneeberger K, Clark RM, Lanz C, Warthmann N, Weigel D: Sequencing of natural strains of Arabidopsis thaliana with short reads. Genome Res 2008, 18:2024–2033.

8. Clark RM, Schweikert G, Toomajian C, Ossowski S, Zeller G, Shinn P, Warthmann N, Hu TT, Fu G, Hinds DA, et al: Common sequence polymorphisms shaping genetic diversity in Arabidopsis thaliana. Science 2007, 317:338–342.

9. Li J, Ding J, Zhang W, Zhang Y, Tang P, Chen JQ, Tian D, Yang S: Unique evolutionary pattern of numbers of gramineous NBS-LRR genes. Mol Genet Genomics 2010, 283:427–438.

10. Meyers BC, Kozik A, Griego A, Kuang H, Michelmore RW: Genome-wide analysis of NBS-LRR-encoding genes in Arabidopsis. Plant Cell 2003, 15:809–834.

11. Zhou T, Wang Y, Chen J-Q, Araki H, Jing Z, Jiang K, Shen J, Tian D: Genome-wide identification of NBS genes in japonica rice reveals significant expansion of divergent non-TIR NBS-LRR genes. Mol Genet Genomics 2004, 271:402–415.

12. Sun X, Cao Y, Yang Z, Xu C, Li X, Wang S, Zhang Q: Xa26, a gene conferring resistance to Xanthomonas oryzae pv. oryzae in rice, encodes an LRR receptor kinase-like protein. Plant J 2004, 37:517–527.

13. Michelmore RW, Meyers BC: Clusters of resistance genes in plants evolve by divergent selection and a birth-and-death process. Genome Res 1998, 8:1113–1130.

14. Hall SA, Allen RL, Baumber RE, Baxter LA, Fisher K, Bittner-Eddy PD, Rose LE, Holub EB, Beynon JL: Maintenance of genetic variation in plants and pathogens involves complex networks of gene-for-gene interactions. Mol Plant Pathol 2009, 10:449–457.

15. Jacob F, Vernaldi S, Maekawa T: Evolution and conservation of plant NLR functions. Front Immunol 2013, 4:297.

16. Schatz MC, Maron LG, Stein JC, Wences AH, Gurtowski J, Biggers E, Lee H, Kramer M, Antoniou E, Ghiban E, et al: Whole genome de novo assemblies of three divergent strains of rice, Oryza sativa, document novel gene space of aus and indica. Genome Biol 2014, 15:506.

17. Yu P, Wang C, Xu Q, Feng Y, Yuan X, Yu H, Wang Y, Tang S, Wei X: Detection of copy number variations in rice using array-based comparative genomic hybridization. BMC Genomics 2011, 12:372.

18. Zheng L-Y, Guo X-S, He B, Sun L-J, Peng Y, Dong S-S, Liu T-F, Jiang S, Ramachandran S, Liu C-M: Genome-wide patterns of genetic variation in sweet and grain sorghum (Sorghum bicolor). Genome Biol 2011, 12:R114.

19. Xu X, Liu X, Ge S, Jensen JD, Hu F, Li X, Dong Y, Gutenkunst RN, Fang L, Huang L, et al: Resequencing 50 accessions of cultivated and wild rice yields markers for identifying agronomically important genes. Nat Biotechnol 2012, 30:105–111.

20. Bush SJ, Castillo-Morales A, Tovar-Corona JM, Chen L, Kover PX, Urrutia AO: Presence–absence variation in A. thaliana is primarily associated with genomic signatures consistent with relaxed selective constraints. Mol Biol Evol 2013, 31:59–69.

21. Kroj T, Chanclud E, Michel-Romiti C, Grand X, Morel JB: Integration of decoy domains derived from protein targets of pathogen effectors into plant immune receptors is widespread. New Phytol 2016, 210:618–626.

22. Sarris PF, Cevik V, Dagdas G, Jones JD, Krasileva KV: Comparative analysis of plant immune receptor architectures uncovers host proteins likely targeted by pathogens. BMC Biol 2016, 14:8.

23. Bailey PC, Schudoma C, Jackson W, Baggs E, Dagdas G, Haerty W, Moscou M, Krasileva KV: Dominant integration locus drives continuous diversification of plant immune receptors with exogenous domain fusions. Genome Biol 2018, 19:23.

24. Triplett LR, Cohen SP, Heffelfinger C, Schmidt CL, Huerta A, Tekete C, Verdier V, Bogdanove AJ, Leach JE: A resistance locus in the American heirloom rice variety Carolina Gold Select is triggered by TAL effectors with diverse predicted targets and is effective against African strains of Xanthomonas oryzae pv. oryzicola. Plant J 2016, 87:472–483.

25. Shields DS (Ed.). The golden seed: writings on the history and culture of Carolina gold rice. Beaufort, South Carolina: Douglas W. Bostick for the Carolina Gold Rice Foundation; 2010.

26. Duitama J, Silva A, Sanabria Y, Cruz DF, Quintero C, Ballen C, Lorieux M, Scheffler B, Farmer A, Torres E, et al: Whole genome sequencing of elite rice cultivars as a comprehensive information resource for marker assisted selection. PLoS One 2015, 10:e0124617.

27. Ayres NM, McClung AM, Larkin PD, Bligh HFJ, Jones CA, Park WD: Microsatellites and a single-nucleotide polymorphism differentiate apparent amylose classes in an extended pedigree of US rice germ plasm. Theor Appl Genet 1997, 94:773–781.

28. Ji Z, Ji C, Liu B, Zou L, Chen G, Yang B: Interfering TAL effectors of Xanthomonas oryzae neutralize R-gene-mediated plant disease resistance. Nat Comm 2016, 7:13435.

29. Read AC, Rinaldi FC, Hutin M, He Y-Q, Triplett LR, Bogdanove AJ: Suppression of Xo1-mediated disease resistance in rice by a truncated, non-DNA-binding TAL effector of Xanthomonas oryzae. Front Plant Sci 2016, 7:1516.

30. Sakaguchi S: Linkage studies on the resistance to bacterial leaf blight, Xanthomonas oryzae (Uyeda et Ishiyama) Dowson, in rice. Bull Natl Inst Agric Sci Ser D 1967, 16:1–18.

31. He Q, Li D, Zhu Y, Tan M, Zhang D, Lin X: Fine mapping of Xa2, a bacterial blight resistance gene in rice. Mol Breed 2006, 17:1–6.

32. Ise K, CY Li, CR Ye, and YQ Sun: Inheritance of resistance to bacterial leaf blight in differential rice variety Asominori. Int Rice Res Notes 1998, 23:13–14.

33. Endo T, Yamaguchi M, Kaji R, Nakagomi K, Kataoka T, Yokogami N, Nakamura T, Ishikawa G, Yonemaru J-i, Nishio T: Close linkage of a blast resistance gene, Pias(t), with a bacterial leaf blight resistance gene, Xa1-as(t), in a rice cultivar ‘Asominori’. Breed Sci 2012, 62:334–339.

34. Ogawa T, Morinaka T, Fujii K, Kimura T: Inheritance of Resistance of Rice Varieties Kogyoku and Java 14 to Bacterial Group V of Xanthomonas oryzae. Jap J Phytopathol 1978, 44:137–141.

35. Taura S, Ogawa T, Tabien R, Khush G, Yoshimura A, Omura T: The specific reaction of Taichung Native 1 to Philippine races of bacterial blight and inheritance of resistance resistance to race 5 (PX0112). Rice Genet Newsl 1987, 4:101–102.

36. Wang C, Wen G, Lin X, Liu X, Zhang D: Identification and fine mapping of the new bacterial blight resistance gene, Xa31(t), in rice. Eur J Plant Pathol 2009, 123:235–240.

37. Cheema KK, Grewal NK, Vikal Y, Sharma R, Lore JS, Das A, Bhatia D, Mahajan R, Gupta V, Bharaj TS, Singh K: A novel bacterial blight resistance gene from Oryza nivara mapped to 38 kb region on chromosome 4L and transferred to Oryza sativa L. Genet Res 2008, 90:397–407.

38. Aravind L: The BED finger, a novel DNA-binding domain in chromatin-boundary-element-binding proteins and transposases. Trends Biochem Sci 2000, 25:421–423.

39. Yoshimura S, Yamanouchi U, Katayose Y, Toki S, Wang Z-X, Kono I, Kurata N, Yano M, Iwata N, Sasaki T: Expression of Xa1, a bacterial blight-resistance gene in rice, is induced by bacterial inoculation. Proc Natl Acad Sci USA 1998, 95:1663–1668.

40. Schornack S, Ballvora A, Gürlebeck D, Peart J, Ganal M, Baker B, Bonas U, Lahaye T: The tomato resistance protein Bs4 is a predicted non-nuclear TIR-NB-LRR protein that mediates defense responses to severely truncated derivatives of AvrBs4 and overexpressed AvrBs3. Plant J 2004, 37:46–60.

41. Kawahara Y, de la Bastide M, Hamilton JP, Kanamori H, McCombie WR, Ouyang S, Schwartz DC, Tanaka T, Wu J, Zhou S: Improvement of the Oryza sativa Nipponbare reference genome using next generation sequence and optical map data. Rice 2013, 6:4.

42. Kolmogorov M, Yuan J, Lin Y, Pevzner PA: Assembly of long, error-prone reads using repeat graphs. Nat Biotechnol 2019, 37:540–546.

43. Zimin AV, Puiu D, Luo MC, Zhu T, Koren S, Marcais G, Yorke JA, Dvorak J, Salzberg SL: Hybrid assembly of the large and highly repetitive genome of Aegilops tauschii, a progenitor of bread wheat, with the MaSuRCA mega-reads algorithm. Genome Res 2017, 27:787–792.

44. Steuernagel B, Jupe F, Witek K, Jones JD, Wulff BB: NLR-parser: rapid annotation of plant NLR complements. Bioinformatics 2015, 31:1665–1667.

45. Bayer PE, Edwards D, Batley J: Bias in resistance gene prediction due to repeat masking. Nat Plants 2018, 4:762–765.

46. Stein JC, Yu Y, Copetti D, Zwickl DJ, Zhang L, Zhang C, Chougule K, Gao D, Iwata A, Goicoechea JL, et al: Genomes of 13 domesticated and wild rice relatives highlight genetic conservation, turnover and innovation across the genus Oryza. Nat Genet 2018, 50:285–296.

47. Wilkins KE, Booher NJ, Wang L, Bogdanove AJ: TAL effectors and activation of predicted host targets distinguish Asian from African strains of the rice pathogen Xanthomonas oryzae pv. oryzicola while strict conservation suggests universal importance of five TAL effectors. Front Plant Sci 2015, 6:536.

48. Chaisson MJ, Tesler G: Mapping single molecule sequencing reads using basic local alignment with successive refinement (BLASR): application and theory. BMC Bioinformatics 2012, 13:238.

49. Bendahmane A, Farnham G, Moffett P, Baulcombe DC: Constitutive gain-of-function mutants in a nucleotide binding site-leucine rich repeat protein encoded at the Rx locus of potato. Plant J 2002, 32:195–204.

50. van Ooijen G, Mayr G, Kasiem MM, Albrecht M, Cornelissen BJ, Takken FL: Structure-function analysis of the NB-ARC domain of plant disease resistance proteins. J Exp Bot 2008, 59:1383–1397.

51. Chen J, Huang Q, Gao D, Wang J, Lang Y, Liu T, Li B, Bai Z, Luis Goicoechea J, Liang C, et al: Whole-genome sequencing of Oryza brachyantha reveals mechanisms underlying Oryza genome evolution. Nat Comm 2013, 4:1595.

52. Wang M, Yu Y, Haberer G, Marri PR, Fan C, Goicoechea JL, Zuccolo A, Song X, Kudrna D, Ammiraju JS, et al: The genome sequence of African rice (Oryza glaberrima) and evidence for independent domestication. Nat Genet 2014, 46:982–988.

53. Leister D: Tandem and segmental gene duplication and recombination in the evolution of plant disease resistance gene. Trends Genet 2004, 20:116–122.

54. Germain H, Seguin A: Innate immunity: has poplar made its BED? New Phytol 2011, 189:678–687.

55. Van de Weyer A-L, Monteiro F, Furzer OJ, Nishimura MT, Cevik V, Witek K, Jones JDG, Dangl JL, Weigel D, Bemm F: The Arabidopsis thaliana pan-NLRome. bioRxiv 2019:537001.

56. Marchal C, Zhang J, Zhang P, Fenwick P, Steuernagel B, Adamski NM, Boyd L, McIntosh R, Wulff BBH, Berry S, et al: BED-domain-containing immune receptors confer diverse resistance spectra to yellow rust. Nat Plants 2018, 4:662–668.

57. Kanzaki H, Yoshida K, Saitoh H, Tamiru M, Terauchi R: Protoplast cell death assay to study Magnaporthe oryzae AVR gene function in rice. Methods Mol Biol 2014, 1127:269–275.

58. Cesari S, Kanzaki H, Fujiwara T, Bernoux M, Chalvon V, Kawano Y, Shimamoto K, Dodds P, Terauchi R, Kroj T: The NB-LRR proteins RGA4 and RGA5 interact functionally and physically to confer disease resistance. EMBO J 2014, 33:1941–1959.

59. Le Roux C, Huet G, Jauneau A, Camborde L, Trémousaygue D, Kraut A, Zhou B, Levaillant M, Adachi H, Yoshioka H, et al: A receptor pair with an integrated decoy converts pathogen disabling of transcription factors to immunity. Cell 2015, 161:1074–1088.

60. Brabham HJ, Hernández-Pinzón I, Holden S, Lorang J, Moscou MJ: An ancient integration in a plant NLR is maintained as a trans-species polymorphism. bioRxiv 2018:239541.

61. Xu X, Chen H, Fujimura T, Kawasaki S: Fine mapping of a strong QTL of field resistance against rice blast, Pikahei-1(t), from upland rice Kahei, utilizing a novel resistance evaluation system in the greenhouse. Theor Appl Genet 2008, 117:997–1008.

62. Xu X, Hayashi N, Wang C-T, Fukuoka S, Kawasaki S, Takatsuji H, Jiang C-J: Rice blast resistance gene Pikahei-1(t), a member of a resistance gene cluster on chromosome 4, encodes a nucleotide-binding site and leucine-rich repeat protein. Mol Breed 2014, 34:691–700.

63. Smith CW: Rice: origin, history, technology, and production. United States of America: John Wiley & Sons; 2002.

64. Yu Y, Tang T, Qian Q, Wang Y, Yan M, Zeng D, Han B, Wu CI, Shi S, Li J: Independent losses of function in a polyphenol oxidase in rice: differentiation in grain discoloration between subspecies and the role of positive selection under domestication. Plant Cell 2008, 20:2946–2959.

65. Jupe F, Witek K, Verweij W, Śliwka J, Pritchard L, Etherington GJ, Maclean D, Cock PJ, Leggett RM, Bryan GJ: Resistance gene enrichment sequencing (RenSeq) enables reannotation of the NB-LRR gene family from sequenced plant genomes and rapid mapping of resistance loci in segregating populations. Plant J 2013, 76:530–544.

66. Witek K, Jupe F, Witek AI, Baker D, Clark MD, Jones JD: Accelerated cloning of a potato late blight-resistance gene using RenSeq and SMRT sequencing. Nat Biotechnol 2016, 34:656–660.

67. Steuernagel B, Periyannan SK, Hernandez-Pinzon I, Witek K, Rouse MN, Yu G, Hatta A, Ayliffe M, Bariana H, Jones JD, et al: Rapid cloning of disease-resistance genes in plants using mutagenesis and sequence capture. Nat Biotechnol 2016, 34:652–655.

68. Stam R, Scheikl D, Tellier A: Pooled enrichment sequencing identifies diversity and evolutionary pressures at NLR resistance genes within a wild tomato population. Genome Biol Evol 2016, 8:1501–1515.

69. Andolfo G, Jupe F, Witek K, Etherington GJ, Ercolano MR, Jones JD: Defining the full tomato NB-LRR resistance gene repertoire using genomic and cDNA RenSeq. BMC Plant Biol 2014, 14:120.

70. Giolai M, Paajanen P, Verweij W, Percival-Alwyn L, Baker D, Witek K, Jupe F, Bryan G, Hein I, Jones J: Targeted capture and sequencing of gene-sized DNA molecules. BioTechniques 2016, 61:315.

71. Arora S, Steuernagel B, Gaurav K, Chandramohan S, Long Y, Matny O, Johnson R, Enk J, Periyannan S, Singh N, et al: Resistance gene cloning from a wild crop relative by sequence capture and association genetics. Nat Biotechnol 2019, 37:139–143.

72. Meyer RS, Choi JY, Sanches M, Plessis A, Flowers JM, Amas J, Dorph K, Barretto A, Gross B, Fuller DQ, et al: Domestication history and geographical adaptation inferred from a SNP map of African rice. Nat Genet 2016, 48:1083.

73. Zimin AV, Puiu D, Hall R, Kingan S, Clavijo BJ, Salzberg SL: The first near-complete assembly of the hexaploid bread wheat genome, Triticum aestivum. Gigascience 2017, 6:1–7.

74. Marcais G, Delcher AL, Phillippy AM, Coston R, Salzberg SL, Zimin A: MUMmer4: A fast and versatile genome alignment system. PLoS Comp Biol 2018, 14:e1005944.

75. Jain M, Koren S, Miga KH, Quick J, Rand AC, Sasani TA, Tyson JR, Beggs AD, Dilthey AT, Fiddes IT, et al: Nanopore sequencing and assembly of a human genome with ultra-long reads. Nat Biotechnol 2018, 36:338–345.

76. Li H, Durbin R: Fast and accurate short read alignment with Burrows-Wheeler transform. Bioinformatics 2009, 25:1754–1760.

77. Garrison EM, Gabor: Haplotype-based variant detection from short-read sequencing. arXiv 2012:1207.3907.

78. Kersey PJ, Allen JE, Allot A, Barba M, Boddu S, Bolt BJ, Carvalho-Silva D, Christensen M, Davis P, Grabmueller C, et al: Ensembl Genomes 2018: an integrated omics infrastructure for non-vertebrate species. Nucleic Acids Res 2018, 46:D802–D808.

79. Wu TD, Watanabe CK: GMAP: a genomic mapping and alignment program for mRNA and EST sequences. Bioinformatics 2005, 21:1859–1875.

80. Kent WJ: BLAT--the BLAST-like alignment tool. Genome Res 2002, 12:656–664.

81. Patro R, Duggal G, Love MI, Irizarry RA, Kingsford C: Salmon provides fast and bias-aware quantification of transcript expression. Nat Methods 2017, 14:417–419.

82. Steuernagel B, Witek K, Krattinger SG, Ramirez-Gonzalez RH, Schoonbeek H-j, Yu G, Baggs E, Witek AI, Yadav I, Krasileva KV, et al: Physical and transcriptional organisation of the bread wheat intracellular immune receptor repertoire. bioRxiv 2018:339424.

83. Quinlan AR, Hall IM: BEDTools: a flexible suite of utilities for comparing genomic features. Bioinformatics 2010, 26:841–842.

84. Chojnacki S, Cowley A, Lee J, Foix A, Lopez R: Programmatic access to bioinformatics tools from EMBL-EBI update: 2017. Nucleic Acids Res 2017, 45:W550–W553.

85. Stamatakis A: RAxML version 8: a tool for phylogenetic analysis and post-analysis of large phylogenies. Bioinformatics 2014, 30:1312–1313.

86. Letunic I, Bork P: Interactive tree of life (iTOL) v3: an online tool for the display and annotation of phylogenetic and other trees. Nucleic Acids Res 2016, 44:W242–W245.

87. Marchler-Bauer A, Bo Y, Han L, He J, Lanczycki CJ, Lu S, Chitsaz F, Derbyshire MK, Geer RC, Gonzales NR, et al: CDD/SPARCLE: functional classification of proteins via subfamily domain architectures. Nucleic Acids Res 2017, 45:D200–D203.

88. Benson G: Tandem repeats finder: a program to analyze DNA sequences. Nucleic Acids Res 1999, 27:573–580.

89. Crooks GE, Hon G, Chandonia JM, Brenner SE: WebLogo: a sequence logo generator. Genome Res 2004, 14:1188–1190.

