## Additional_file_12_PPO_figure for "Genome assembly and characterization of a complex zfBED-NLR gene-containing disease resistance locus in Carolina Gold Select rice with Nanopore sequencing"

Polymorphism at the polyphenol oxidase gene linked to BB and blast resistance genes at the Xo1 locus

a)

```
CGS_nonfunctional_PPO    1 MESINVAPGTTATPRMAPPPPPPCITNLQSTLRYNNLLLHRKTKGWKPRNVSCRVDRRDV  60
                           MESINVAPGTTATPRMAPPPPPPCITNLQSTLRYNNLLLHRKTKGWKPRNVSCRVDRRDV
MH63_functional_PPO      1 MESINVAPGTTATPRMAPPPPPPCITNLQSTLRYNNLLLHRKTKGWKPRNVSCRVDRRDV  60

CGS_nonfunctional_PPO   61 LLGISGAAAMVATQGGGGALAAPIQAPDLGDCHQPVDVPATAPAINCCPTYSAGTVAVDF 120
                           LLGISGAAAMVATQGGGGALAAPIQAPDLGDCHQPVDVPATAPAINCCPTYSAGTVAVDF
MH63_functional_PPO     61 LLGISGAAAMVATQGGGGALAAPIQAPDLGDCHQPVDVPATAPAINCCPTYSAGTVAVDF 120

CGS_nonfunctional_PPO  121 APPPASSPLRVRPAAHLADRAYLAKYERAVSLMKKLPADDPRSFEQQWRVHCAYCDGAYD 180
                           APPPASSPLRVRPAAHLADRAYLAKYERAVSLMKKLPADDPRSFEQQWRVHCAYCDGAYD
MH63_functional_PPO    121 APPPASSPLRVRPAAHLADRAYLAKYERAVSLMKKLPADDPRSFEQQWRVHCAYCDGAYD 180

CGS_nonfunctional_PPO  181 QVGFPGLEIQIHSCWLFFPWHRMYLYFHERILGKLIGDETFALPFWNWDAPDGMSFPAIY 240
                           QVGFPGLEIQIHSCWLFFPWHRMYLYFHERILGKLIGDETFALPFWNWDAPDGMSFPA+Y
MH63_functional_PPO    181 QVGFPGLEIQIHSCWLFFPWHRMYLYFHERILGKLIGDETFALPFWNWDAPDGMSFPAMY 240

CGS_nonfunctional_PPO  241 ANRWSPLYDPRRNQAHLPPFPLDLDYSGTDTNIPKDQLIDQNLNIMYRQMISGARKAELF 300
                           ANRWSPLYDPRRNQAHLPPFPLDLDYSGTDTNIPKDQLIDQNLNIMYRQMISGARKAELF
MH63_functional_PPO    241 ANRWSPLYDPRRNQAHLPPFPLDLDYSGTDTNIPKDQLIDQNLNIMYRQMISGARKAELF 300

CGS_nonfunctional_PPO  301 MGQPYRAGDQPEPGAGTVESVPHNPVHRWTGDPRQPNGEDMGIFYSAARDPVFFAHHGNV 360
                           MGQPYRAGDQPEPGAGTVESVPHNPVHRWTGDPRQPNGEDMGIFYSAARDPVFFAHHGNV
MH63_functional_PPO    301 MGQPYRAGDQPEPGAGTVESVPHNPVHRWTGDPRQPNGEDMGIFYSAARDPVFFAHHGNV 360

CGS_nonfunctional_PPO  361 DRMWHIR----------LHRPRLARRQLLLLRRGGPPRPRPRPGHPRPVGAALHVPGRGS 410
                           DRMWHIR             P                          P          G
MH63_functional_PPO    361 DRMWHIRRGLLFPGDTDFTDPDWLDASFFFYDEEARLVRVRVRDTLDPSALRFTYQDVGL 420

CGS_nonfunctional_PPO  411 PVAERQAVHGSSQHAGARGRRVPGD-----------PGQDRAGGRDEA-------QGVEE 452
                           P    +   G++          P             P   R+    E        +G+E
MH63_functional_PPO    421 PWLNAKPSTGAASTPAPAAGAFPATLDKTVRVAVTRPRASRSREEKEEEEEVLVIEGIEI 480

CGS_nonfunctional_PPO  453 P----------------RGEGGGGGGARHRGDRDPRPLHVRQVRRVRERARERGRRG--D 494
                           P                 G+G     A   G     P  + +   ++ R  E  R G  D
MH63_functional_PPO    481 PDHSTYVKFDVFVNAPESGDGAATCAATCAGSVALAPHGIHREGQLSPRKTE-ARFGICD 539

CGS_nonfunctional_PPO  495 VRGDVRRQRRAGAARDPPRGAAVAEEDGGEVRHMRLAGRHRRRRRQDDRRVDRAEVWLRF 554
                           +  D+      GA  D     ++    G +   + +AG                   +
MH63_functional_PPO    540 LLDDI------GADGDKTIVVSIVPRCGCD--SVTVAG-------------------VSI 572

CGS_nonfunctional_PPO  555 GHRRRRQHRLR 565
                           G+ +
MH63_functional_PPO    573 GYAK------- 576
```

b)

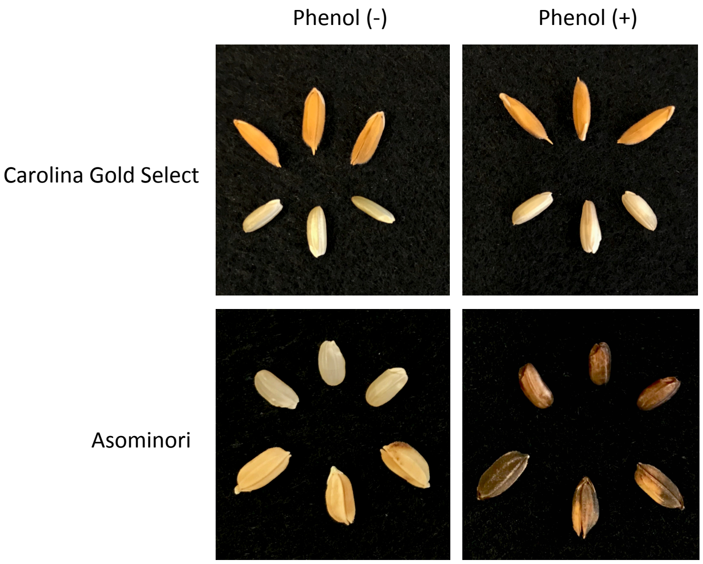


(a) Amino acid sequence alignment of *Oryza sativa* indica cultivar MH63 functional polyphenol oxidase (PPO) gene with the sequence of Carolina Gold Select PPO. Loss-of-function mutation highlighted in yellow. (b) Polyphenol oxidase activity in seeds of the known PPO positive rice cultivar Asominori and Carolina Gold Select with and without hull. Dark brown indicates activity.
