## Additional_file_11_zfBED_nt_tree for "Genome assembly and characterization of a complex zfBED-NLR gene-containing disease resistance locus in Carolina Gold Select rice with Nanopore sequencing"

Additional file 11
Nucleotide based maximum likelihood zfBED domain tree

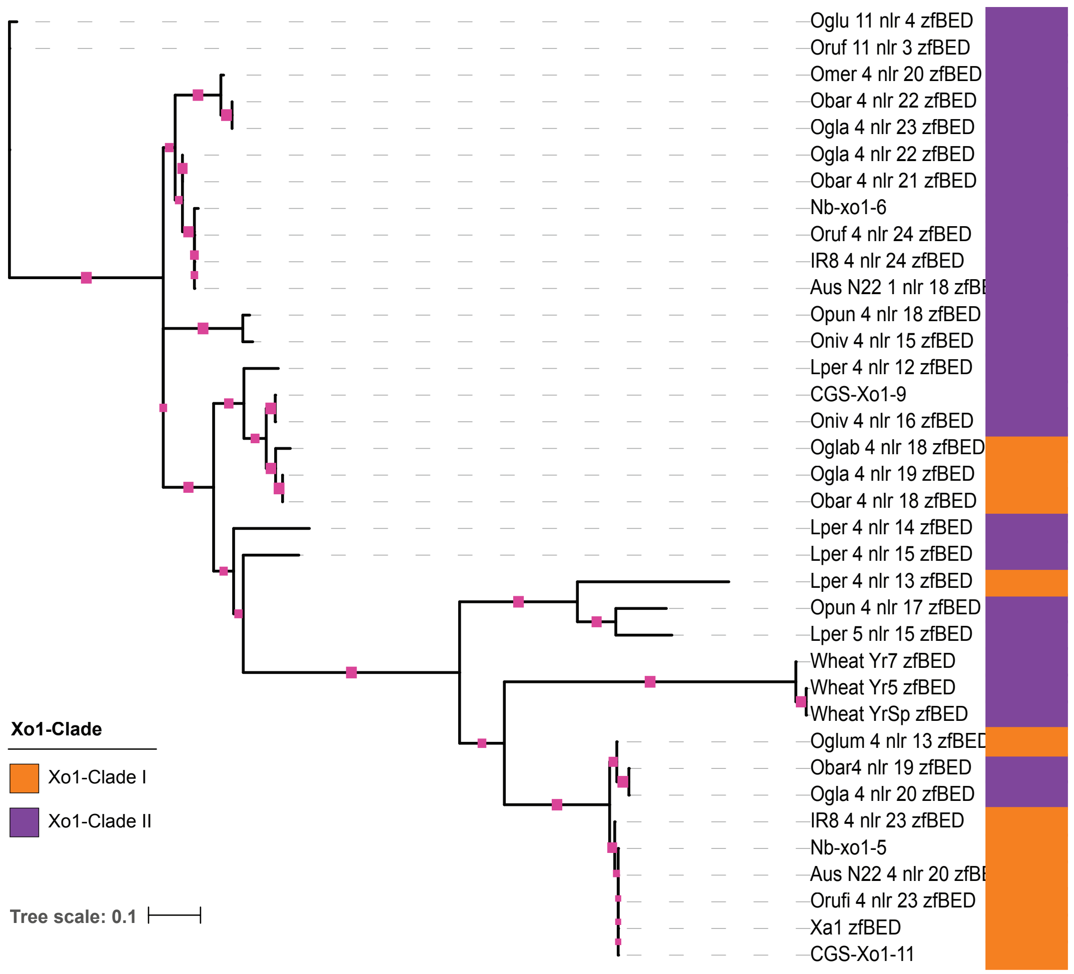


Maximum likelihood tree based on nucleotide sequences of zfBED domains from Xo1 clade I and II NLRs identified across Oryzeae. Pink squares indicate bootstrap support >80. Tree file is available at iTOL – http://itol.embl.de/shared/acr242
