## Additional_file_10_BEDsequences for "Genome assembly and characterization of a complex zfBED-NLR gene-containing disease resistance locus in Carolina Gold Select rice with Nanopore sequencing"

**Additional file 10 – NLR zfBED domain nucleotide sequences**

>Wheat_Yr7

TCCCCGGTATGGGAACACTTCACCATCACAGAAACAACTATCGACGGGAAGCGTTCAAAAGCCAAATGTAAGTACTGTGGAAATGATTTTAATTGCGAAACGAAGACAAACGGGACTTCATCTATGAAAAAACATTTGGAGAAGGAGCATTCC

>Wheat_Yr5

TCCCCGGTATGGGAACACTTCACGATCACAGAAACAACTATCGACGGGAAGCGTTCAAAAGCCAAATGTAACTACTGTGGAAATGATTTTAATTGCGAAACGAAGACAAACGGGACTTCATCTATGAAAAAACATTTGGAGAAAGAGCATTCC

>Wheat_YrSp

TCCCCGGTATGGGAACACTTCACGATCACAGAAACAACTATCGACGGGAAGCGTTCAAAAGCCAAATGTAACTACTGTGGAAATGATTTTAATTGCGAAACGAAGACAAACGGGACTTCATCTATGAAAAAACATTTGGAGAAAGAGCATTCC

>CGS_4_nlr_17

GTATGGCAGCACTTTGAGATTGTGGAAGAGAAAAACGGTAAGCCCGCGAAAGCCAAGTGTATCGACTGTCACACGGTGGTCAAGTGCGGCTCCGACAATGGGACATCGGTTTTGCACAATCACCGCAACAGTGGCAAATGTAAAAGG

>Xa1

AAGGCATGGGAACACTTTACTACCGTAGAGTTCACTGCTGACGGGAAGGATTCTAAAGCACGGTGCAAGTACTGCCACAAGGACCTATGTTGCACATCTAAGAACGGGACATCAGCTTTGCGCAACCATCTCAATGTTTGCAAGAGG

>Aus_N22__4_nlr_20_zfBED

AAGGCATGGGAACACTTTACTACCGTAGAGTTCACTGCTGACGGGAAGGATTCTAAAGCACGGTGCAAGTACTGCCACAAGGACCTATGTTGCACATCTAAGAACGGGACATCAGCTTTGCGCAACCATCTCAAT

>IR8__4_nlr_23_zfBED

AAGGCATGGGAACACTTTACTACCGTAGAGTTCACTGCTGACGGGAAGGATTCTAAAGCACGGTGCAAGTACTGCCACAAGGACCTATGTTGCACATCTAAGAACGGGACATCAGCTTTGCGCAACCATCTCAATGTTTACAAGAGGAAACGT

>Lper_4_nlr_13_zfBED

TTTGTATGGCAACACTTTGTCAGAATAAAAGATGCGAACGGGAAACTCGTGCAAGCAAGGTGCAAGTACTGTCACAAGACGCTGAAATGCCCAACCGAAAACGGGACAAATTCTTTGAGGAGACATCACTATAGCAAA

>CGS_4_nlr_19_zfBED

AAGGCATGGGAACACTTTACTACCGTAGAGTTCACTGCTGACGGGAAGGATTCTAAAGCACGGTGCAAGTACTGCCACAAGGACCTATGTTGCACATCTAAGAACGGGACATCAGCTTTGCGCAACCATCTCAAT

>Nippo_4_nlr_20_zfBED

AAGGCATGGGAACACTTTACTACCGTAGAGTTCACTGCTGACGGGAAGGATTCTAAAGCACGGTGCAAGTACTGCCACAAGGACCTATGTTGCACATCTAAGAACGGGACATCAGCTTTGCGCAACCATCTCAAT

>Obarthii_4_nlr_18_zfBED

GTATGGCAATACTTTGAGATCGTGGAAGAGAAAAACGGAAAGCCTGCGAAAGCCAAGTGTGTCGACTGTCACACGGTGGTCAAGTGCGGCTCCGACAATGGGACTTCGGTTTTGCACAATCACCGCAACAGTGGCAAATGTAAAAGG

>Oglab_4_nlr_18_zfBED

GTATGGCAGCACTTTGAGATCGTGGAAGAGAAAAACGGAAAGCCTGCGAAAGCCAAGTGTGTCGACTGTGGCACGGTGGTCAAGTGCGGCTCCGACAATGGGACTTCGGTACTGCACAATCACCGCAACAGTGGCAAATGTAAAAGG

>Oglum_4_nlr_13_zfBED

AAGGCATGGGAACACTTTACTCCCGTAGAGTTCACTGCTGATGGGAAGGCTTCTAAAGCACGGTGCAAGTACTGCCACAAGGACCTATGTTGCACATCTAAGAACGGGACATCAGCTTTGCGCAACCATCTCAAT

>Orufi_4_nlr_23_zfBED

AAGGCATGGGAACACTTTACTACCGTAGAGTTCACTGCTGACGGGAAGGATTCTAAAGCACGGTGCAAGTACTGCCACAAGGACCTATGTTGCACATCTAAGAACGGGACATCAGCTTTGCGCAACCATCTCAAT

>Ogla_4_nlr_19

GTATGGCAATACTTTGAGATCGTGGAAGAGAAAAACGGAAAGCCTGCGAAAGCCAAGTGTGTCGACTGTCACACGGTGGTCAAGTGCGGCTCCGACAATGGGACTTCGGTTTTGCACAATCACCGCAACAGTGGCAAATGTAAAAGG

>Lper_5_nlr_15_zfBED

TGGAAAAAGTTCACCAGAATACCAAATAAAAAGGGAAAGATCGAGAGAGCAAGGTGCAACTACTGCAACAAGGAGCTGATGTGCCCATCCAAAAATGGGACAAGTACTTTGCGCAATCATCCTAAGAGT

>Oglu_11_nlr_4_zfBED

GTGGTATGGAAGAACTTTGATATCACTGAATATGGAAATGGAAAGGCTGTGAAGAAGGTAAAATGTATTCACTGTAACACTGTGCTCAAGTGTGGTGCTAGCAAAGGGACGTCGGTTTTGCATAAGCACCTCCGTAGCATC

>Lper_4_nlr_12_zfBED

GAGGTATGGCAGCACTTTGATATTGTGGATGAGAAGAACGGAAAGCCTGCGAAAGCCAAATGTATTGACTGTCATAAGGTGCTCAAGTGTGGTTCCGACAATGGGACATCGGTTTTGCACAATCAT

>Opun_4_nlr_17_zfBED

GTGTGGAACGACTTCTACAGAATACTGGATGAAAAGGGAAAGATCGTGAAAGCAAGGTGCAACTACTGCCAGAAGGAGCTGAAATGCCCATCCAACAATGGGACAAGTATTCTGCGCAATCATCTTAATTGTAAA

>Opun_4_nlr_18_zfBED

TCCAAGGCATGGGCGCACTTCAAAGAGGAAAATGGAAAGCCTGGGAAGGCAAGATGTATTCACTGTCACACGGTGGTCAAGTGCGGTTCTGACAAAGGGACATCGGTTTTGCATAATCACCTCAAGAGT

>Oniv_4_nlr_15_zfBEDSuperfamily

TCCGAGGCATGGGCGCACTTTAAAGAGGAAAATGGAAAGCCTGGGAAGGCAAGATGTATTCACTGTCACACGGTGGTCAAGTGCAGTTCTGACAAAGGGACATCTGTTTTGCATAATCACCTCAAGAGT

>Aus_N22_1_nlr_18_zfBED

AAGGCATGGGGGCACTTTGATATCACTGAAGAAGAAAATGGAAAGCCTGTGAAGGCAAGGTGTATTCACTGTCACACGGTGGTCAAGTGCGGTTCTGAAAAAGGGACATCGGTTTTGCATAATCACCTCAAGAGT

>Nippo_4_nlr_21_zfBED

AAGGCATGGGGGCACTTTGATATCACTGAAGAAGAAAATGGAAAGCCTGTGAAGGCAAGGTGTATTCACTGTCACACGGTGGTCAAGTGCGGTTCTGAAAAAGGGACATCAGTTTTGCATAATCACCTCAAGAGT

>Oruf_4_nlr_24_zfBED

AAGGCATGGGGGCACTTTGATATCACTGAAGAAGAAAATGGAAAGCCTGTGAAGGCAAGGTGTATTCACTGTCACACGGTGGTCAAGTGCGGTTCTGAAAAAGGGACATCGGTTTTGCATAATCACCTCAAGAGT

>IR8__4_nlr_24_zfBED

AAGGCATGGGGGCACTTTGATATCACTGAAGAAGAAAATGGAAAGCCTGTGAAGGCAAGGTGTATTCACTGTCACACGGTGGTCAAGTGCGGTTCTGAAAAAGGGACATCGGTTTTGCATAATCACCTCAAGAGT

>Obar_4_nlr_22_zfBED

AAGGCTTGGGGGCACTTCGACATCACCGAAGAGGAGAATGGGAAGCCACTGAAGGCAAGATGTATTCACTGTCACACGGTGGTCAGGTGTACTTCTGACAAAGGGACATCGGTTTTGCACAACCATCTCAAGAGTGACAGTTGCAAAAAG

>Ogla_4_nlr_23_zfBED

AAGGCTTGGGGGCACTTCGACATCACCGAAGAGGAGAATGGGAAGCCACTGAAGGCAAGATGTATTCACTGTCACACGGTGGTCAGGTGTACTTCTGACAAAGGGACATCGGTTTTGCACAACCATCTCAAGAGTGACAGTTGCAAAAAG

>Omer_4_nlr_20_zfBED

AAGGCATGGGGACACTTCGACATCACCGAAGAGGAGAATGGGAAGCCACTGAAGGCAAGATGTATTCACTGTCACACGGTGGTCAAGTGTACTTCTGACAAAGGGACATCGGTTTTGCACAACCACCTCAAGAGTGACAGTTGCAAAAAG

>Obar_4_nlr_21_zfBED

AAGGCATGGGGGCACTTCGATATCACTGAAGAAGAAAATGGAAAGCCTGTGAAGGCAAGGTGTATTCACTGTCACACGGTGGTCAAGTGCGGTTCTGACAAAGGGACATCGGTTTTGCACAATCACCTCAAGAGT

>Ogla_4_nlr_22_zfBED

AAGGCATGGGGGCACTTCGATATCACTGAAGAAGAAAATGGAAAGCCTGTGAAGGCAAGGTGTATTCACTGTCACACGGTGGTCAAGTGCGGTTCTGACAAAGGGACATCGGTTTTGCACAATCACCTCAAGAGT

>Lper_4_nlr_15_zfBED

GAGGTATGGCGGCACTTCACGGTCGTGGAAAGGGAAAATGGGAAGACTGTGAAAGCCAGATGTATTGACTGCAATACGGTGGTCAAGTGCGGTTCCAGCAATGGAACATCAGTTTTGCACAATCAC

>Obar4_nlr_19_zfBED

GCATGGGAACACTTTACTCCCGTAGAGTTCACTGCTGGTGGGAAGGCTTCTAAAGCACGATGCAAGTACTGCGACAAGGACCTATGTTGCACATCTAAGAACGGGACATCAGCTTTGCGCAACCATCTCAAT

>Ogla_4_nlr_20_zfBED

GCATGGGAACACTTTACTCCCGTAGAGTTCACTGCTGGTGGGAAGGCTTCTAAAGCACGATGCAAGTACTGCGACAAGGACCTATGTTGCACATCTAAGAACGGGACATCAGCTTTGCGCAACCATCTCAAT

>Oniv_4_nlr_16_zfBED

GTATGGCAGCACTTTGAGATTGTGGAAGAGAAAAACGGTAAGCCCGCGAAAGCCAAGTGTATCGACTGTCACACGGTGGTCAAGTGCGGCTCCGACAATGGGACATCGGTTTTGCACAATCACCGCAACAGTGGCAAATGTAAAAGG

>Oruf_11_nlr_3_zfBED

GAGGTATGGAAGAACTTTGATATCACTGAACATGGAAATGGAAAGGCTGTGAAGAAGGTAAAATGTATTCACTGTAACACTGTGCTCAAGTGTGGTGCTAGCAAAGGGACGTCGGTTTTGCATAAGCACCTCCGTAGCATC

>Lper_4_nlr_14_zfBED

AAAGTTTGGAATCACTTCAATATCGAGGAAGAGGAAAATGGGAAGGCTACGTTATCAAGATGTATTGACTGTCATACGGTGGTCAAGTGCGGTTCAAAAAATGGGACATCTGTTTTGCACAATCACCGCAAGAGTAAA
