## Additional_file_8_Maximum_likelihood_tree_Oryzeae for "Genome assembly and characterization of a complex zfBED-NLR gene-containing disease resistance locus in Carolina Gold Select rice with Nanopore sequencing"

Maximum likelihood tree of NLR genes across Oryzeae

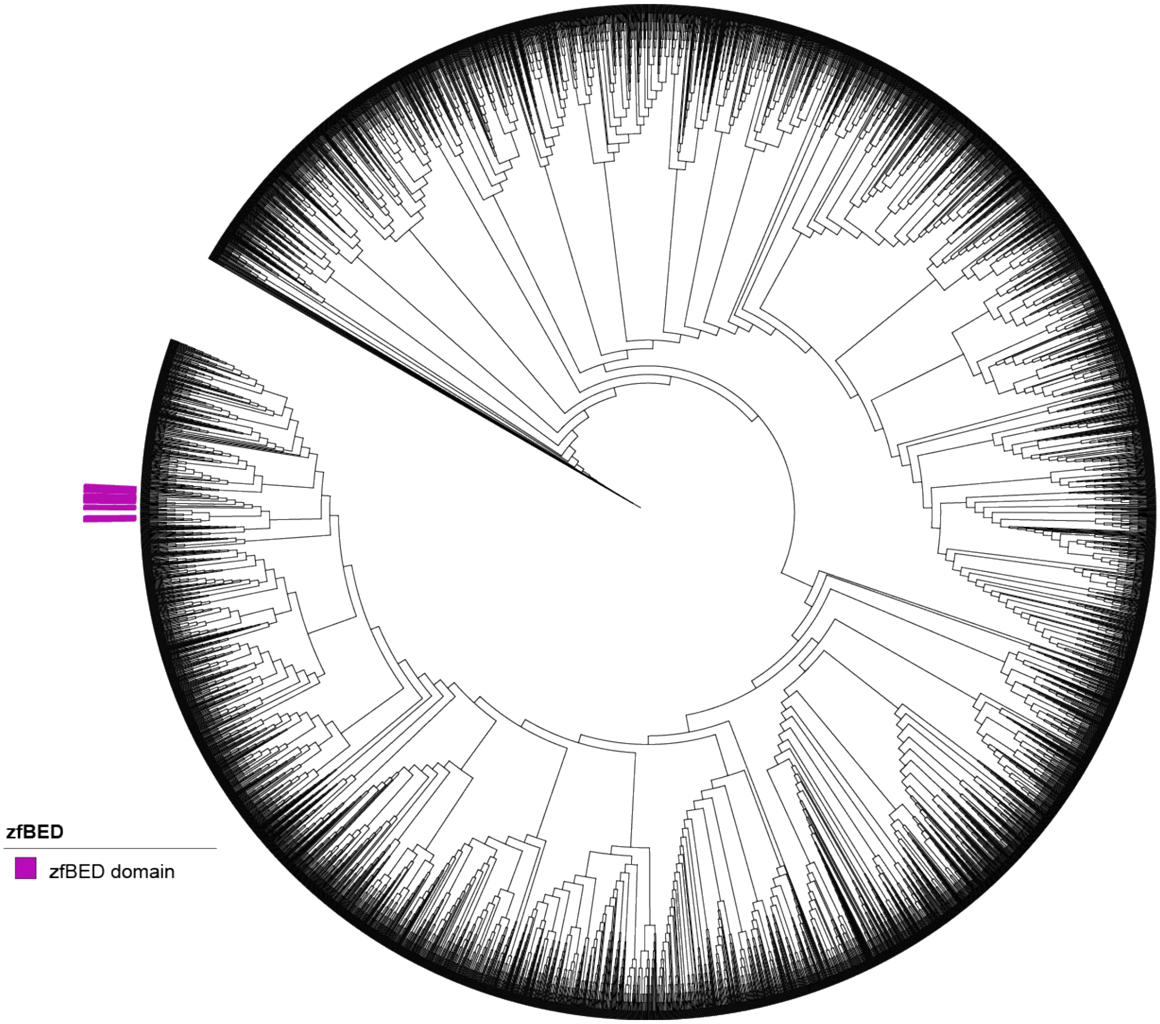


Maximum likelihood tree of 5104 NB-ARC domain amino acid sequences detected by NLR-Annotator in representative Oryzeae genomes. Tree includes known rice R-genes (as in Additional file 3) and three wheat zfBED-NLRs. Xo1 clade zfBED NLRs are annotated on the tree. NB-ARC amino acid sequences are available in Additional file 3. Tree file is available at iTOL – http://itol.embl.de/shared/acr242
