## Additional_file_7_TandemRptAlignment for "Genome assembly and characterization of a complex zfBED-NLR gene-containing disease resistance locus in Carolina Gold Select rice with Nanopore sequencing"

Tandem Repeats in CGS-Xo1$_{11}$, Xa1, and Nb-xo1$_5$

a)

| aa position | *CGS-Xo1$_{11}$* | *Xa1* | *Nb-xo1$_5$* | |
|---|---|---|---|---|
| 8 | L | **W** | **W** | Exon 1 |
| 31 | E | **E** | **E** | |
| 56 | V | **V** | **V** | |
| 72 | S | **S** | **S** | |
| 426 | K | **N** | **N** | NB-ARC |
| 431 | **S** | S | **S** | |
| 549 | **S** | **S** | C | |
| 627 | **M** | **M** | V | "MHV" |
| 755 | **T** | **T** | T | |
| 851 | **Q** | **Q** | Q | |
| 1056 | **S** | **S** | C | |
| 1079 | Y | **C** | **C** | |
| Repeats start | | | | |

b)

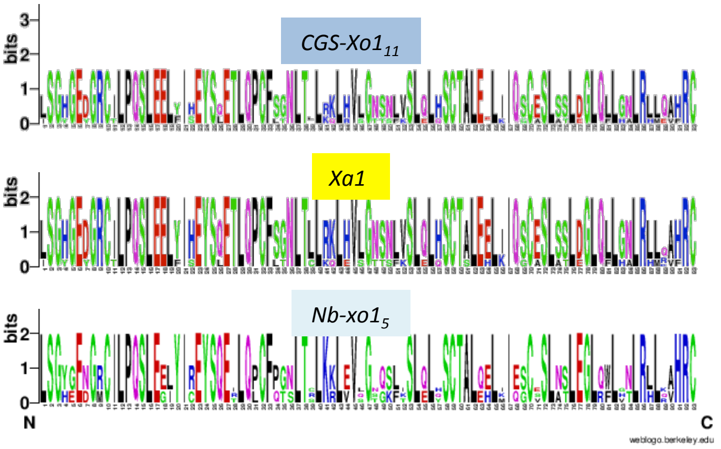


c)

| | Repeat # | *CGS-Xo1$_{11}$* 1 | 2 | 3 | 4 | Final | *Xa1* 1 | 2 | 3 | 4 | 5 | Final | *Nb-xo1$_5$* 1 | 2 | 3 |
|---|---|---|---|---|---|---|---|---|---|---|---|---|---|---|---|
| *CGS-Xo1$_{11}$* | 1 | **100** | **97.9** | **97.9** | **95** | **73.1** | 99.6 | 97.9 | 98.2 | 98.2 | 97.9 | 73.8 | 84.6 | 85.3 | 71.7 |
| | 2 | **97.9** | **100** | **98.6** | **95.7** | **74.6** | 97.5 | 100 | 98.9 | 98.9 | 98.6 | 75.3 | 86 | 85.7 | 72 |
| | 3 | **97.9** | **98.6** | **100** | **96.4** | **75.3** | 97.5 | 98.6 | 98.9 | 98.9 | 98.6 | 76 | 86.7 | 86.4 | 72.4 |
| | 4 | **95** | **95.7** | **96.4** | **100** | **77.1** | 94.6 | 95.7 | 96.1 | 96.1 | 95.7 | 77.8 | 85.3 | 86 | 72.8 |
| | Final | **73.1** | **74.6** | **75.3** | **77.1** | **100** | 72.8 | 74.6 | 74.9 | 74.6 | 74.6 | 98.6 | 72.8 | 75.3 | 87.5 |
| *Xa1* | 1 | 99.6 | 97.5 | 97.5 | 94.6 | 72.8 | **100** | **97.5** | **97.9** | **97.9** | **97.5** | **73.5** | 84.2 | 85.7 | 71.3 |
| | 2 | 97.9 | 100 | 98.6 | 95.7 | 74.6 | **97.5** | **100** | **98.9** | **98.9** | **98.6** | **75.3** | 86 | 85.7 | 72 |
| | 3 | 98.2 | 98.9 | 98.9 | 96.1 | 74.9 | **97.9** | **98.9** | **100** | **99.3** | **98.9** | **74.9** | 85.7 | 85.3 | 72.4 |
| | 4 | 98.2 | 98.9 | 98.9 | 96.1 | 74.6 | **97.9** | **98.9** | **99.3** | **100** | **98.9** | **75.3** | 85.7 | 85.3 | 72 |
| | 5 | 97.9 | 98.6 | 98.6 | 95.7 | 74.6 | **97.5** | **98.6** | **98.9** | **98.9** | **100** | **75.3** | 85.3 | 85 | 72 |
| | Final | 73.8 | 75.3 | 76 | 77.8 | 98.6 | **73.5** | **75.3** | **74.9** | **75.3** | **75.3** | **100** | 73.5 | 76.3 | 88.2 |
| *Nb-xo1$_5$* | 1 | 84.6 | 86 | 86.7 | 85.3 | 72.8 | 84.2 | 86 | 85.7 | 85.7 | 85.3 | 73.5 | **100** | **87.5** | **70.3** |
| | 2 | 85.3 | 85.7 | 86.4 | 86 | 75.3 | 85.7 | 85.7 | 85.3 | 85.3 | 85 | 76.3 | **87.5** | **100** | **76** |
| | 3 | 71.7 | 72 | 72.4 | 72.8 | 87.5 | 71.3 | 72 | 72.4 | 72 | 72 | 88.2 | **70.3** | **76** | **100** |

Percent Identity

60 80 100

(a) List of all DNA polymorphisms and resulting amino acid identity upstream of the tandem repeat region when comparing the three predicted coding sequences. Bold amino acid codes indicate a silent mutation, final column indicates protein motif or region. (b) WebLogos showing amino acid conservation of the repeat units. (c) Heatmap of repeat unit percent identity within and among the three coding sequences. *Nb-xo1$_5$* encodes an additional cryptic final repeat that does not align and is not included in (b) or (c).
