## Additional_file_5_IR8vsCGS_dotplot for "Genome assembly and characterization of a complex zfBED-NLR gene-containing disease resistance locus in Carolina Gold Select rice with Nanopore sequencing"

Additional file 5
Dotplot comparison of the Xo1 locus in Carolina Gold Select and indica cultivar IR8

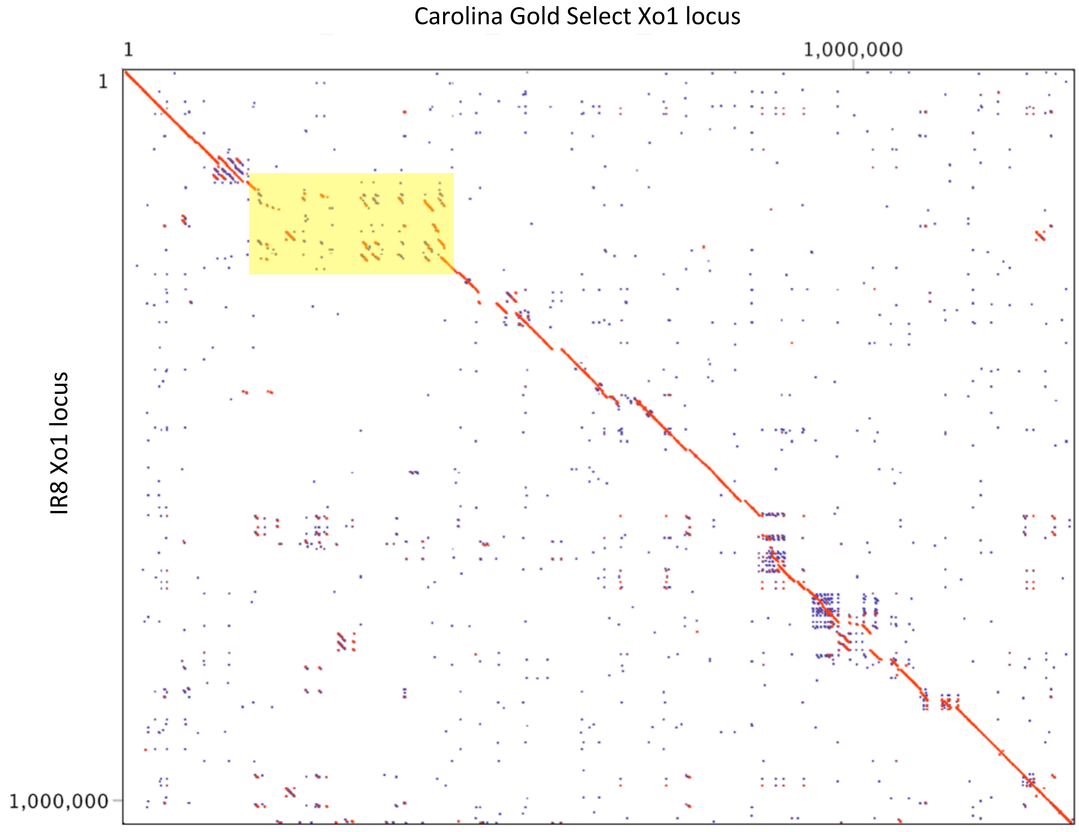


Dotplot comparison of the Xo1 region of Carolina Gold Select (chr4 22729801..24027920) and indica cultivar IR8 (chr4 31657596..32688920). The yellow box highlights the insertion that is present in Carolina Gold Select and absent in IR8.
